# Variation in the life history strategy of cells underlies tumor’s functional diversity

**DOI:** 10.1101/829135

**Authors:** Tao Li, Jialin Liu, Jing Feng, Zhenzhen Liu, Sixue Liu, Minjie Zhang, Yuezheng Zhang, Yali Hou, Dafei Wu, Chunyan Li, Young-Bin Chen, Chung-I Wu, Hua Chen, Xuemei Lu

## Abstract

Classical *r*- vs. *K*-selection theory describes the trade-offs between high reproductive output and competitiveness and guides research in evolutionary ecology^1–5^. While its impact has waned in the recent past, cancer evolution may rekindle it^6–10^. Indeed, solid tumors are an ideal theater for *r*- and *K*-selection and, hence, a good testing ground for ideas on life-history strategy evolution^11,12^. In this study, we impose *r*- or *K*-selection on HeLa cells to obtain strongly proliferative r cells and highly competitive K cells. RNA-seq analysis indicates that phenotypic trade-offs in r and K cells are associated with distinct patterns of expression of genes involved in the cell cycle, adhesion, apoptosis, and contact inhibition. Both empirical observations and simulations based on an ecological competition model show that the trade-off between cell proliferation and competitiveness can evolve adaptively and rapidly in naïve cell lines. It is conceivable that the contrasting selective pressure may operate in a realistic ecological setting of actual tumors. When the r and K cells are mixed *in vitro*, they exhibit strikingly different spatial and temporal distributions in the resultant cultures. Thanks to this niche separation, the fitness of the entire tumor increases. Our analyses of life-history trade-offs are pertinent to evolutionary ecology as well as cancer biology.

## Introduction

Diverse environmental conditions act on populations and species, leading to selection-driven emergence of niche-specific adaptive phenotypes and preventing the emergence of a “superorganism”^13^. Such a superorganism, often dubbed “Darwinian demon,” would produce very large numbers of offspring and live indefinitely^14^. Existence of such entities is contrary to life history theory and empirical observation. Indeed, evolution of adaptive traits is typically restricted by fitness constrains^15^. These constrains often take the form of trade-offs whereby a life history trait can affect different components of fitness in opposite directions. Thus, directional evolution of such a trait would increase some measures of organismal performance at the expense of others^16^. The trade-offs are fundamentally shaped by the way the organism allocates its energy and resources between reproduction and survival^17,18^. Due to the complexity of life-history traits and environmental variables, empirical measurement of plausible trade-offs and their driving forces remains difficult^15^.

In contrast to natural organisms, cancers appear to be exempt from all constraints during the process of somatic cell evolution. A series of biological features, the so-called “hallmarks of cancer”, are characterized by fast proliferation, resistance to low oxygen and crowded environment, and the ability to recruit blood vessels and escape the immune system^19^. How can all aspects of fitness be maximized in cancers? Perhaps heterogeneity within tumors enables several cell lineages to adopt a variety of characteristics and colonize different niches in a changing environment^12,20–25^. The internal and external microenvironments that cancer cells are confronted with in a multicellular organism are akin to complex ecosystems^25–34^. Trade-offs between cell proliferation and survival may apply to such cancer cell populations^25,33,35^. Both rapid cell proliferation and stable survival strategies must complement each other to achieve high fitness of a tumor as a whole^12^. Therefore, cancer cell populations can be used to test selection pressures and adaptive strategies that govern the trade-off between increasing proliferation and survival, and the ecological mechanisms that underlie these trade-offs in heterogenous populations.

An important and well-defined environmental variable governing evolutionary change is population density relative to essential resources^36^. The theory of density-dependent natural selection, often called *r*- and *K*-selection, states that at extreme population densities evolution produces alternative strategies^37^. The trade-offs are presumed to arise because the genotypes with the highest fitness at high population densities have low fitness at low density and vice-versa^15,38^. The r-populations are selected for high intrinsic rate of growth (r) in environments where population density is low and resources are abundant but perform badly at high density. In contrast, K-populations, experiencing strong competition for limited resources under high density conditions, should evolve high intraspecific competitive ability and enhance their carrying capacity (K). K-selected populations do not have high growth rates because they are near the carrying capacity for their environment^1,2,5,39,40^.

In this study, we performed artificial selection for cell density on HeLa cell lines in order to amplify the diversity of cell growth within tumors. We asked whether selection under different density regimes modifies per capita growth rates and competitiveness as predicted by models that postulate a trade-off between *r-* and *K*-selection. To examine the phenotypic trade-offs at the molecular level, we carry out RNA-seq and explore the specific gene expression and pathway characteristics of r and K cells. The dynamics of density-dependent population growth in mixed populations change with the proportions of r and K cells within them. We model these dynamics and fit our models to empirical observations in order to quantify the interaction among the various trade-off phenotypes in a heterogenous population and their effect on fitness of whole tumors.

## Results

### Density-dependent selection and fitness changes of r- and K-selected cell populations

The initial cell population (*IN* cells) was a single cell clone from a HeLa cell line. When the size of the population reached 10^7^ cells, we divided the clone in two sub-populations. One sub-population was marked with eGFP (IN_G) and the other with dsRed (IN_R) through lentivirus transfection. Three *r*-selection replicates (using IN_G cells) and three *K*-selection replicates (using IN_R cells) were derived independently. After approximately 200 passages under *r*-selection (the low-density condition) and about 130 passages under *K*-selection (the high-density condition), we obtained six populations of r-selected (r cells) and six of K-selected cells (K cells). The density-dependent selection scheme is illustrated in Figure 1a.

**Figure 1.**
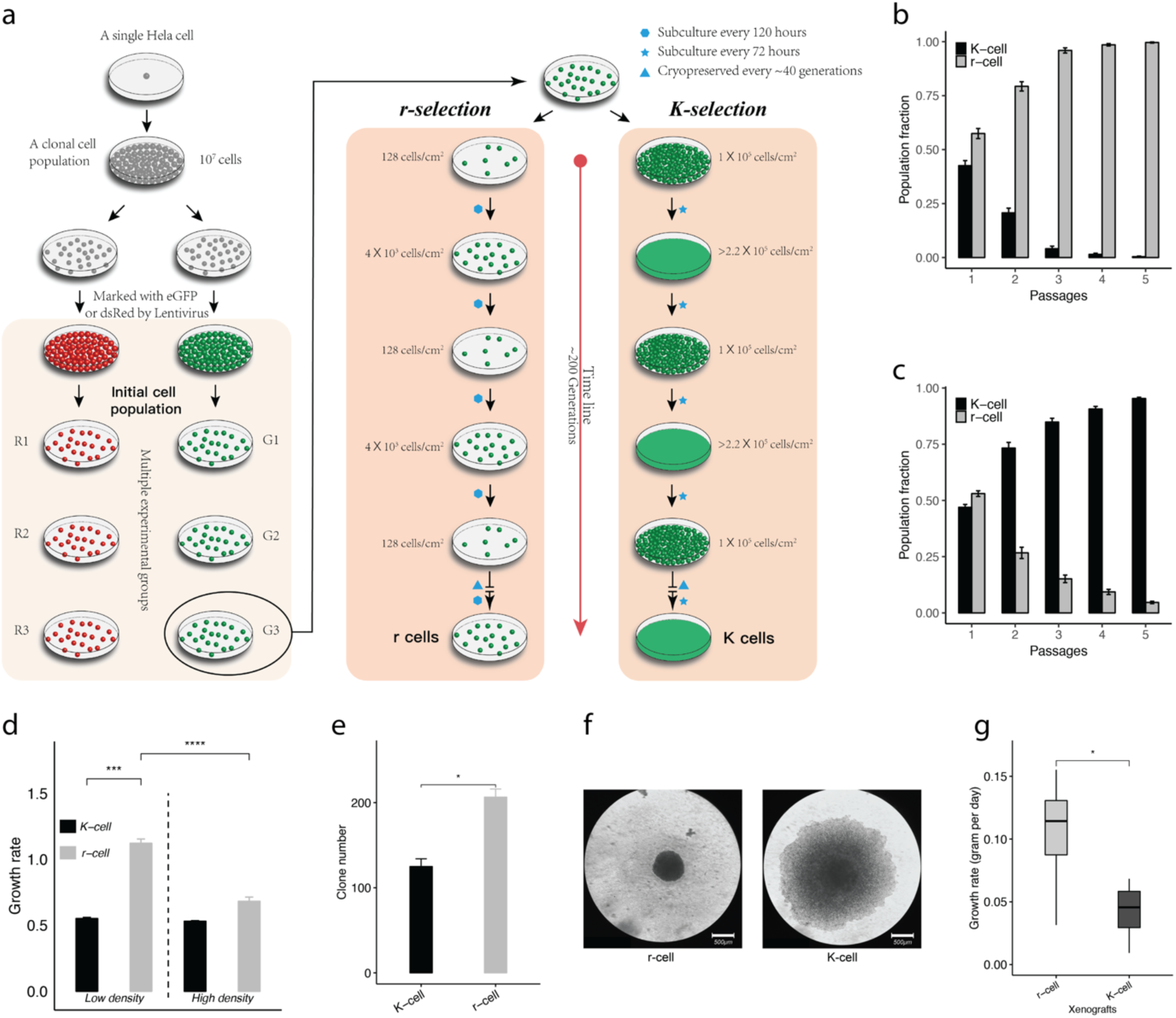
*r*- and *K*-selection in HeLa cells and their growth rate cells in 2D and 3D cultures. **a) *r*- and *K*-selection strategies**. An initial single cell clone was split into six populations, with three labeled with dsRed (R; red dots) and three with eGFP (G; green dots). Each cell culture was passaged >200 times at low (*r*-selection) and high (*K*-selection) density. **Fitness tests were performed at b) low and c) high density**. The Y-axis is the proportion of r and K cells estimated by flow cytometry during five passages (x-axis) of r-K mixed cell cultures. **d) The growth rate of r and K cells across culture conditions**. Cells in 111 r- and 141 K-cell clones were counted every 24 hours. Growth rate is calculated based on cell number change within seven days. **The tumorigenicity of r and K cells is presented based on the number e) and size f) of tumor clones in a soft agar assay on the 7th 21st day. g) The growth rate of r- and K-cell xenografts.** Error bars represent standard deviations. Dash lines separate culture conditions or strategies. Error bars represent standard deviations. Student’s t-test: *P<0.05, **P<0.01, ***P<0.005, ****P<0.0001. Scale bars in **f)** represent 500μm. n= 3 independent experiments per population in b), c), d), and e); n= 12 xenografts in g). mean ± SD.

To test whether the selected r and K cells are more adapted to their corresponding conditions than the ancestral IN cells, we pairwise co-cultured the three types of cells at high and low density. *r* cells become dominant within two passages (three days, Extended Data Figure 1a) in the r-IN mix, suggesting that the r cells have evolved higher fitness than IN cells under these conditions. Likewise, K cells rapidly take over the K-IN mixed population (in four days, two passages, Extended Data Figure 1b). Both r and K cells display better fitness than their counterpart in the r-K mix under corresponding selection conditions (Figure 1b and 1c). We thus successfully selected for alternative life histories in our experiment.

### Density-dependent rates of population growth of r and K cells in 2D- and 3D-growth environments

To explore the possibility that the r and K cells exhibit a trade-off in their density-dependent population growth, we first measured the growth rates of these cells in 2D *in vitro* systems at low and high density. Under low-density, r cells grow faster than K cells (Figure 1d). When the test was performed at high density, there is no significant difference between r and K cells, whereas growth rates of r-cell populations decrease remarkably compared to low density conditions (Figure 1d).

We next tested the difference between r and K cells in their density-dependent rates of population growth in 3D cellular environments. We quantified tumorigenicity by measuring colony growth and formation in a semi-solid agarose gel. r cells displayed a significantly higher rate of colony formation than K cells within seven days (Figure 1e). However, K-cell clones were significantly larger than r-cell colonies on day 21 (Figure 1f). The diameter of K-cell clones was 5 mm on average, while it was 0.5 mm for r-clones. This suggests K-cells have evolved to tolerate high density better than r-cells.

Xenograft mouse models were used to investigate the population growth rate of r and K cells in vivo. Cells were injected into the inguinal skin of BALB/c Nude mice. Tumor nodules were established in all xenografts in about two weeks. The mean growth rate of r-cell tumors was significantly higher than that of K-cell tumors in vivo (Figure 1g).

### Trade-off between cell proliferation and survival in r- and K-cells

The net rate of population growth is determined by both cell death and birth rates. Using annexin-V and DAPI staining, reflecting cell death and the G0/G1 phase of the cell cycle, we measured the proportions of dead cells and distinguished the resting/quiescent (G0/G1) from total cells in the r and K populations at high and low density. Figure 2a shows that the proportion of G0/G1 phase cells is lower in the r-than in the K-cell populations, indicating that r cells proliferate relatively quickly at both low and high density. It also demonstrates that K cell birth rate does not increase at high density.

**Figure 2.**
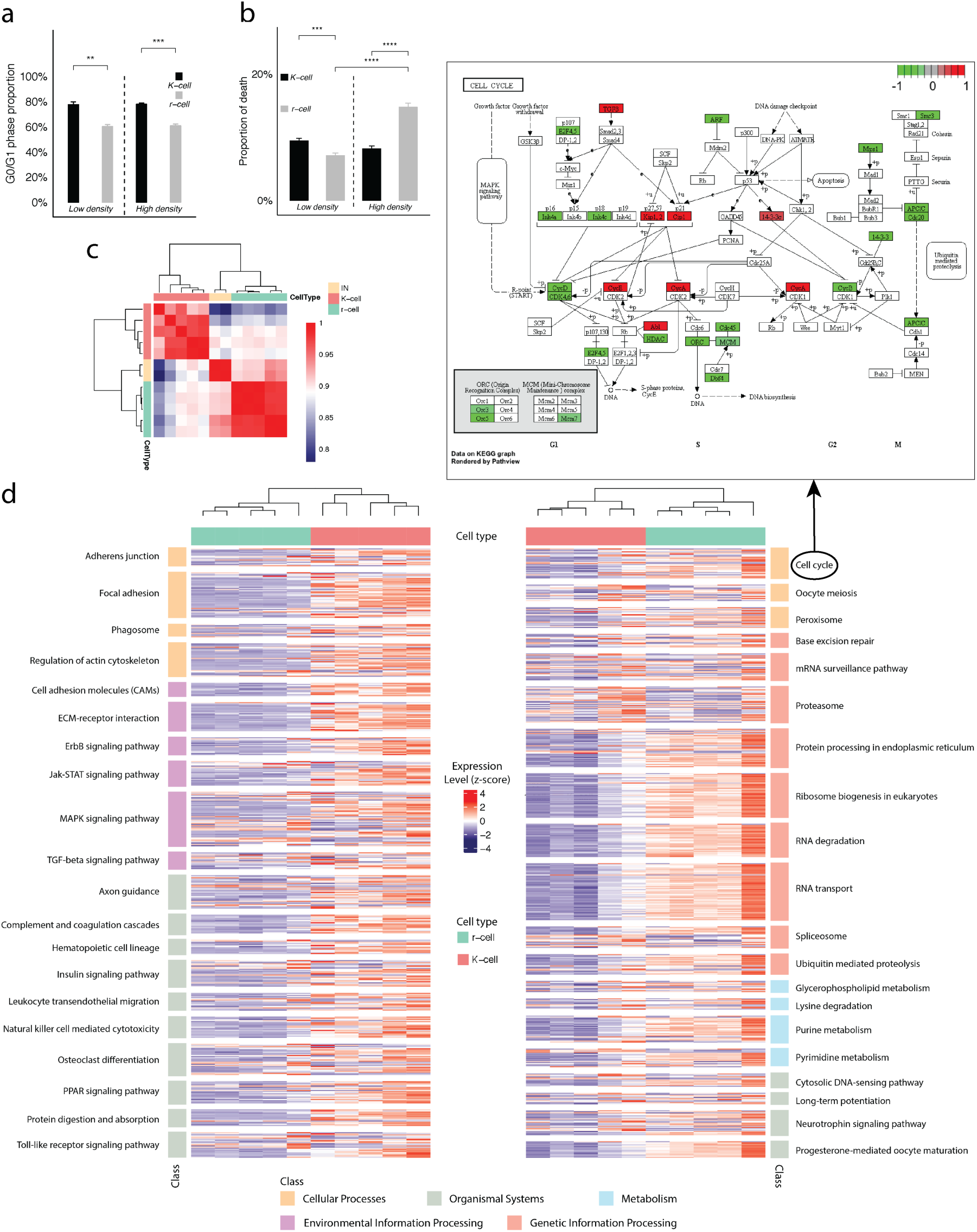
Differences in cell cycle, cell death and gene expression between *r* and *K* cells. **a) The G0/G1 phase proportion and b) the proportion of cell death** in r (gray) and K cells (black) are analyzed using PI and Annexin V staining via flow cytometry under high- and low-density conditions. Dashed lines separate culture conditions. Error bars represent standard deviations. Student’s t-test: *P<0.05, **P<0.01, ***P<0.005, ****P<0.0001. n=3 independent experiments per population**. c) Gene expression correlation between IN, r-, and K-cell populations. d) Pathways that show significantly different expression between r and K cells.** The left panel presents signaling pathways that are overexpressed in five K-cell populations (pink), the right presents pathways overexpressed in five r-cell populations (green). The z-score heatmap indicates the scale of gene expression difference. The upper panel shows the cell cycle pathway with relatively over-(red) and under-(green) expressed genes in r vs. K cells highlighted.

The *K* cell death rate is relatively stable under both conditions (Figure 2b). In contrast, the *r* cell death rate increases significantly under high compared to low density. The *r* cells also die more frequently at high density than *K* cells (Figure 2b). The high birth and death rates of *r* cells suggest that they have evolved to quickly produce offspring rather than to increase their survival, while *K* cells tend to ensure offspring quality rather than number. The high incidence of cell death leads to a decrease in growth rate of *r* cells at high density, and the effect of density in *r-*selected populations is mainly on cell death.

### Transcriptome characteristics support a trade-off between cell proliferation and survival in r- and K-cells

To find molecular characteristics that may be correlated with the phenotypic trade-offs in r and K cells, we carried out RNA-seq in 22 samples, including two replicates of initial cell populations, five K cell lines, five r-cell lines under routine cell culture conditions, and r- and K-replicate lines under high density stress. Multiple comparisons were performed among transcriptional profiles of cell lines across and within density conditions. Differentially expressed genes in these comparisons were identified using standard methods^41,42^. Figure 2c shows that r and IN cell populations cluster closely and differ from the K-cell populations under routine cell-culture conditions (at low density). We detect that 3161 genes show significant difference in gene expression (DEGs) between the r and K cells (with 1748 up- and 1413 down-regulated genes in K cells, Extended Data Table 1). Using the Functional Annotation Tool from the DAVID package, we found 25 pathways significantly enriched for these differentially expressed genes. The top three of these are the spliceosome, pathways involved in cancer, and ribosome biogenesis (Extended Data Table 2).

Genes from the same signaling pathway may increase or decrease its overall expression level, resulting in the enhancement or inhibition of related biological functions^43–45^. The top 20 highly expressed pathways in K or r cells based on the GAGE (General Applicable Gene-set Enrichment for Pathway Analysis)^46^ are listed in Figure 2d. The upregulated pathways in K cells include cell and focal adhesion, ErbB signaling, ECM-receptor interaction, phagosome, regulation of actin cytoskeleton, and Jak-STAT signaling. The cell cycle (upper panel in Figure 2d), metabolism, and genetic information processing (such as ribosome biogenesis and mRNA surveillance) pathways are significantly highly expressed in r cells (Figure 2d).

We next detected the transcriptional difference in responding to density constraints between r and K cells. Dramatic change at the transcriptional level is found in r cells when they are grown at high density. The expression levels of 6373 genes are significantly different from low density (Extended Data Table 1), while the number of DEGs is 2278 in K cells (Extended Data Table 1). Compared to the gene expression profiles under low-density conditions, 1775 genes (859 genes up-regulated; 916 genes down-regulated) present the same trend of expression change in both r and K cells under high density. These are involved in metabolic and serial RNA related pathways. These results suggest that high culture density has a prominent effect on cell metabolism (Extended Data Table 3). In addition to these common changes, only 503 (=2278-1775) genes respond to density change specifically in K cells. The number of genes (6373-1775=4598 genes) responding to the density change in r cells is approximately nine times larger than that, indicating that r cells are more sensitive and less stable at high density than K cells.

### Underrepresentation of contact inhibition in K cells

The direct cellular response to cell density is contact inhibition which mediates cell growth and proliferation via interplay between growth signaling pathways and density constraints. Contact inhibition of proliferation is typically absent in cancer cells^47^. Both RNA-seq analysis and trypsinization assay showed that K cells are prone to form cell-cell adhesion at high density (Figure 2d and Extended Data Figure 6), implying a loss or decrease of contact inhibition^48^. In contrast, cell cycle arrest and the slower growth may still be triggered in r cells by signaling pathways that downregulate proliferation in a cell-density dependent manner^49^. One such pathway is the Hippo-YAP signaling pathway, which is largely responsible for inhibiting cell growth and controls organ size in many organisms^50^. The RNA_seq results in this study show that expression of YAP/TAZ is significantly upregulated in K cells, while the hippo-signaling pathway is overrepresented in gene expression comparison between r and K cells (Extended Data Table 2). In addition, the crosstalk among the hippo signaling and eight other pathways (including adherens junction, focal adhesion, tight junction, PI3K-Akt signaling, mTOR signaling, ErbB signaling, TGF-beta signaling, and Wnt signaling) constructs a regulation network associated with cell cycle, cell survival, cell proliferation, and apoptosis^51–53^. A gene cluster analysis shows that the r and K cells can be distinguished by the expression profile of DEGs involved in these nine signaling pathways (Extended Data Figure 2).

The expression of anti-apoptotic factors can be activated by the transport of dephosphorylated YAP into the cell nucleus^52^. In reacting to high cell density, activated LATS1/2 regulates phosphorylation of the coactivator YAP/TAZ, promoting cytoplasmic localization of YAP and leading to cell apoptosis and restriction of organ overgrowth^54^. Overexpression or hyperactivation of YAP/TAZ has been observed in many types of tumors, stimulating growth and proliferation^55–57^. We performed an immunofluorescence assay to identify the localization of YAP/TAZ in r and K cells under both low- and high-density conditions. The localization of YAP/TAZ in the cytoplasm and nuclei was observed in both r and K cells at low density (Extended Data Figure 3a). In contrast, the nuclear localization of YAP/TAZ is absent in r cells but is still maintained in K cells grown at high density (Figure 3a). This suggests that YAP/TAZ phosphorylation is inhibited in K cells under high density, resulting in the loss of cell contact inhibition^58^. Consequently, cell apoptosis may be triggered by cytoplasmic localization of YAP in r cells but not in K cells as cell density increases.

**Figure 3.**
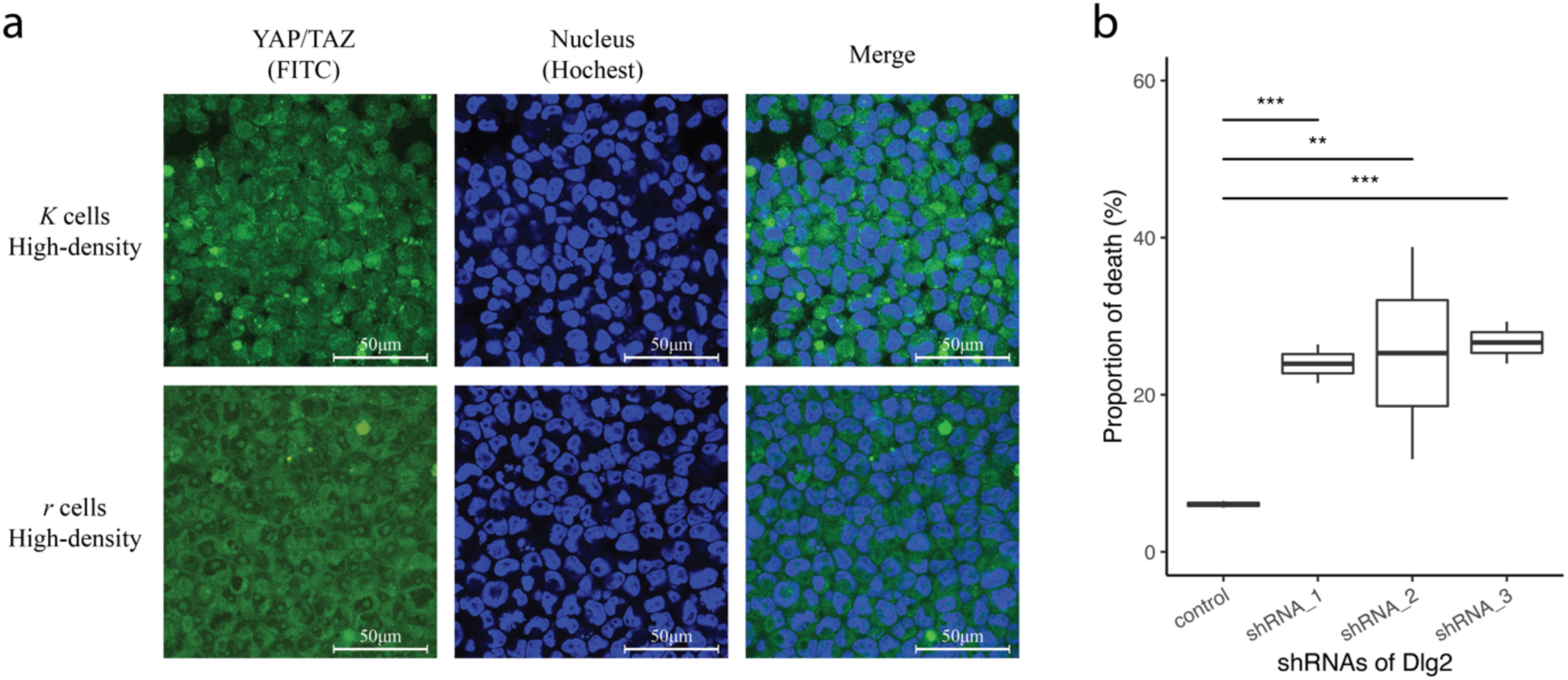
YAP/TAZ colocalization in *r* and *K* cells under high density andthe effect of Dlg-2 knock-down in K cells. **a) YAP/TAZ colocalization in the cytoplasm and nuclei under high density.** YAP/TAZ was immunofluorescently stained with FITC. Hoechst staining marks nuclei. Scale bars represent 50μm. **b) The proportion of cell death in Dlg-2 knockdown *K* cells under high density.** The death rate was measured by Annexin V staining via flow cytometry. Student’s *t*-test: *P<0.05, **P<0.01, ***P<0.005. n=8 independent experiments per population mean ± SD.

In addition, Dlg-2 is a cell polarity gene in the hippo signaling pathway, regulating the inhibition of phosphorylated active YAP/TAZ proteins in the cytoplasm^51,59^. Our transcriptome analysis shows that expression of Dlg-2 is significantly higher in K cells at high than at low density (Extended Data document 1). We confirmed this by RT-PCR (Extended Data Figure 3b). We carried out an siRNA assay to knock down the expression of Dlg-2 in K cells (Extended Data Figure 3c). The apoptosis rate of Dlg-2 knock-down K cells significantly increased at high density (Figure 3b), confirming that the high expression level of Dlg-2 contributes to survival of K cells grown under these conditions.

### Dynamics of density-dependent population growth and competitiveness of r and K cells

#### 1) Empirical observations

Population proportion changes, as well birth and death rates of r and K cells were measured in a co-culture assay. When r and K cells are co-cultured at high density, the proportion of r cells decreases over time (Figure 1c, Extended Data Figure 3) and the death incidence of r cells is significantly higher than of K cells (Figure 4a). The death rate and G0/G1 phase proportion among r cells in co-cultures are both significantly higher than when the r cells are cultured individually under crowded conditions (Figure 4a and 4b). Compared to r, K cells have a relatively stable incidence of death and proportion of cells in G0/G1 phase under co-culture or in individual cultures, although their death rate increases under co-culture (Figure 4a and 4b). These results show that the birth of r cells is restrained and cell death is accelerated when these two different types of cells are cultured together at high density, suggesting that they are in competition when they coexist.

**Figure 4.**
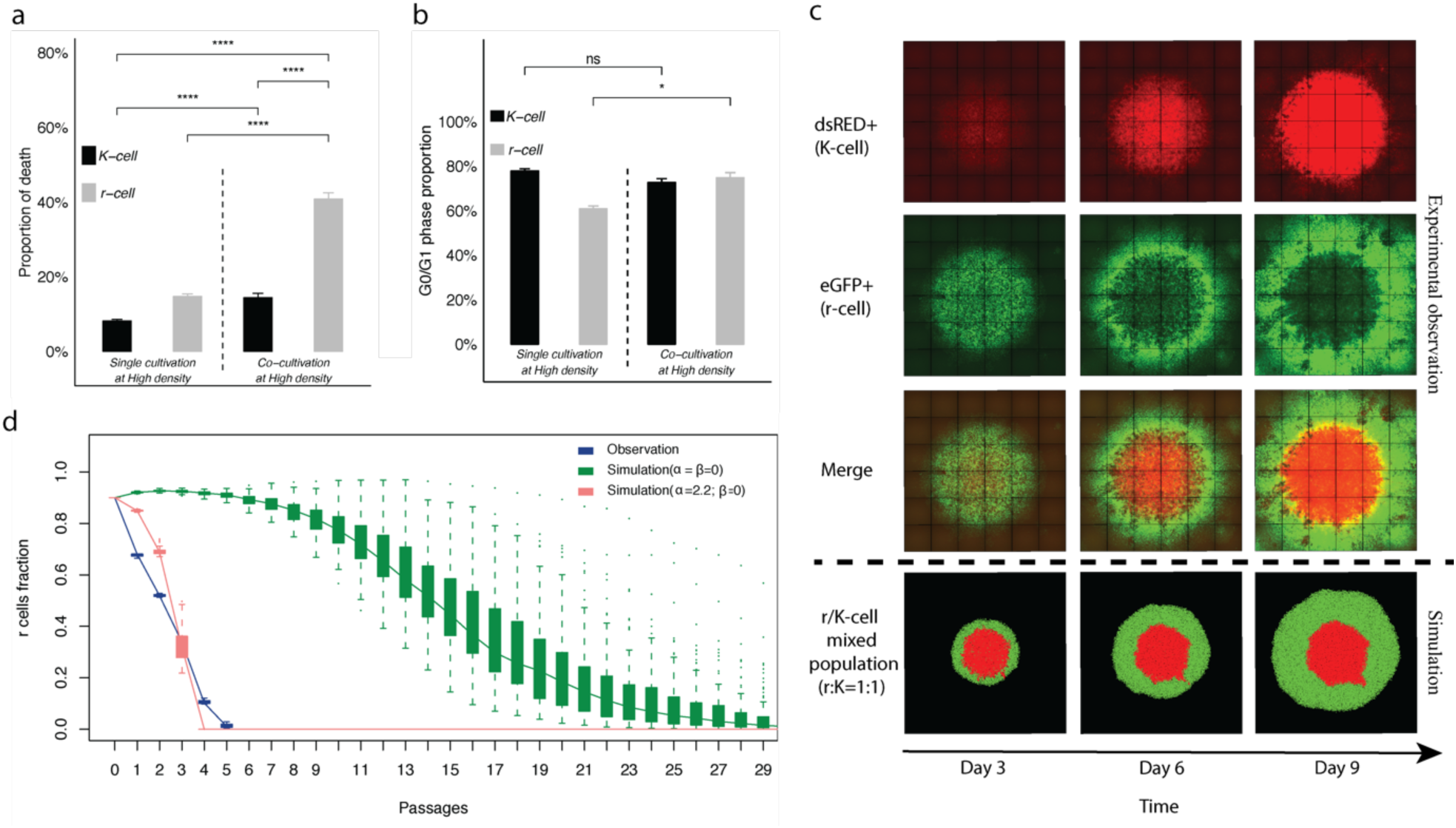
Inter-population interaction and temporal and spatial growth of r and K cells in mixed populations. **a) Cell death and b) G0/G1 phase proportion of r and K cells in individual and mixed cultures.** The y-axis in a) and b) shows death rates and G0/G1 phase proportion of r (gray) and K (black) cells. Death rates were measured by Annexin V staining. G0/G1 phase proportions were measured by PI staining via flow cytometry. Cells were cultured alone or co-cultivated at high density. Dashed lines separate culture strategies. Error bars represent standard deviations. Student’s t-test: ns: non-significant, *P<0.05, ****P<0.0001. n=3 independent experiments per population. **c) Spatial structure in an r-K mixed population.** K and r cells are well mixed in equal proportion and seeded in the center of a six-well plate with total cell number ~10^6^. Each column represents time points from day 3 to day 9 after cell seeding. r and K cells are eGFP and dsRed positive shown in green and red, respectively. The top and bottom panels show the spatial distribution of r and K cells in empirical observations and computer simulations, respectively. **d) The distribution of r cell fractions estimated in vitro (blue) and in silico (red (**α=2.2, β=0**) and green (**α=β=0)). The co-culture of r and K cells is initiated with r/K ratio of 9:1. The y-axis reflects the fraction of r cells in the co-culture; x-axis represents cell passages. (n=100 stochastic simulations per population; n=3 independent experiments; mean ± SD).

Competition may result in niche separation among co-existing populations in an ecological community^60^. To examine this possibility, we carried out co-cultures where approximately 10^6^ r and K cells were well mixed at equal proportion and seeded in the centers of wells in six-well plates. Three replicate co-cultures were scanned every 72 hours. We observed that r and K cells in the co-culture assay tended to occupy different regions in a well. The r cells disperse to the periphery, while K cells grow and occupy the crowded central area (Figure 4c). This observation reveals an additional density-dependent difference in the phenotypes of r and K cells^61,62^.

#### 2) Simulation and parameter estimation

To investigate the inter-population relationship between r and K cells, we adopt the Lotka-Volterra model which has been widely used to study population interaction^63,64–54^. Mixed populations were initiated in our computer simulations with different fractions of r and K cells (Materials and Methods), followed by 30 cell passages at high density. We compared the growth curves of r and K populations in the simulation to the empirical observations described in the previous section. Figure 4d shows that even when the initial proportion of r cells was highest (r:K=9:1) the extinction time of r cells in the simulation with no between-cell type interaction (α = β =0; no effect of one cell population on the other) is approximately five times longer than observed. Simulations reveal that the extinction time of the r cell population is shortened when α is higher than β (Extended Data Figure 5). Comparing the growth curves from empirical observations (blue line in Figure 4d) and in simulations across values of α and β (green and red lines in Figure 4d), we find that the values of α = 2.2 and β = 0 fit the data best (Figure 4d, Extended Data Table 4). Thus, we infer that there is an interaction between r and K cells and K cells influence r cell death.

### Phenotypic diversity promotes cancer cell population growth

#### In silico

To test whether the existence of phenotypic diversity and inter-population interaction promote total fitness, we first carried out stochastic simulations to compare the growth dynamics of r-K mixed populations to pure r- and K-cell assemblages. Unlike in the previous section, the current computational model considers space and density heterogeneity in the environment where the tumor cells grow, and the interaction of r and K cells in these conditions. The rates of cell division and death depend on local cell density. Due to the density effect, cells are able to divide and migrate only if there is sufficient nearby space. The simulation is described in detail in the Materials and Methods and Extended Data Figure 13. Figure 4c illustrates that in silico growth distribution of r and K cells in the mixed population is consistent with empirical observations (the upper panel of Figure 4c). Among-cell interaction and the density effect promote the re-localization of r and K cells, from well-mixed at the beginning of cell culture to a biased distribution with the entire occupation of the K cells in the middle and the outward spread of r cells (the bottom panel of Figure 4c).The mixed populations exhibit significantly higher rate of growth than the pure r- or K-cell populations (Figure 5a and 5b).

**Figure 5.**
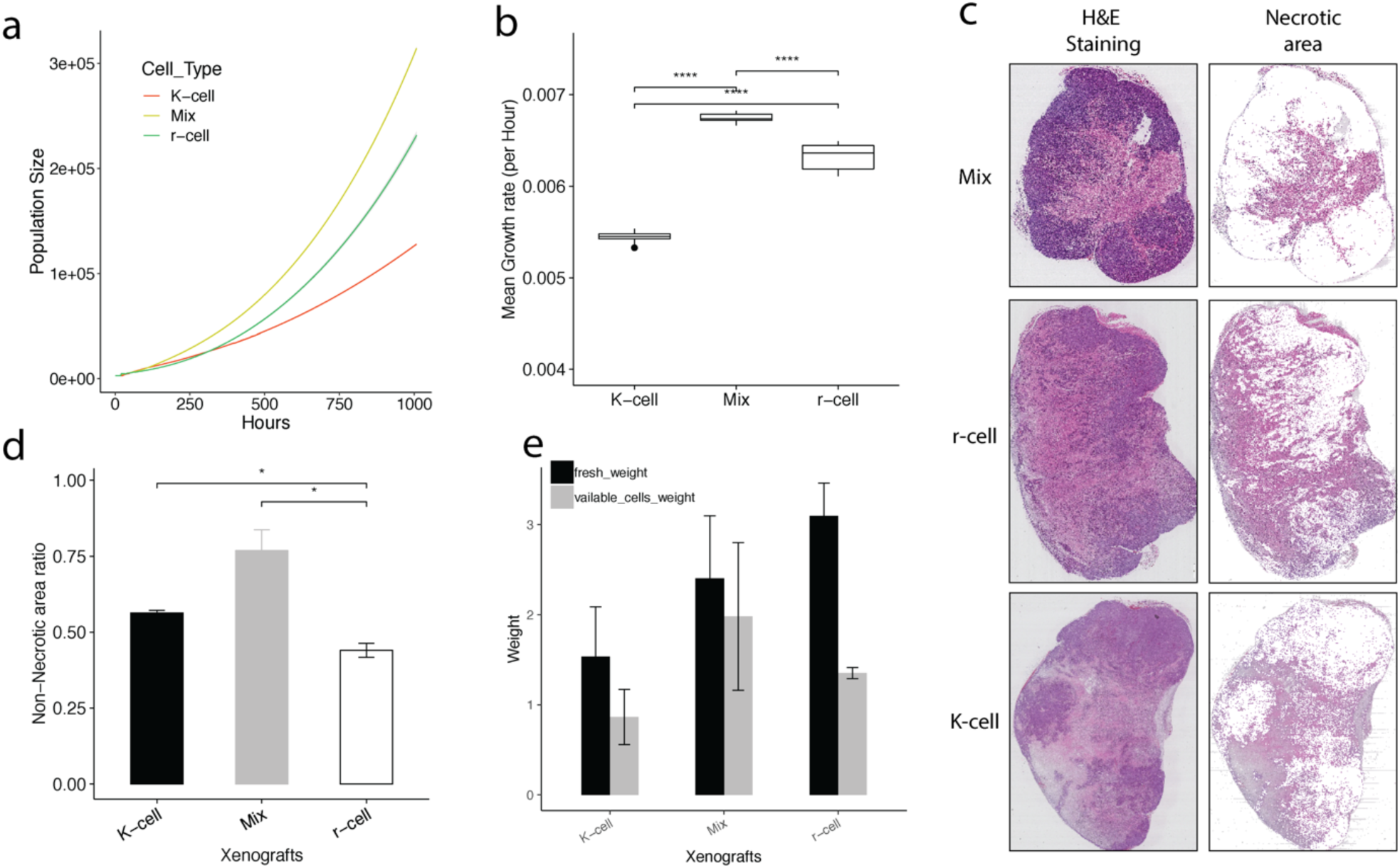
Populartion fitness of r, K, and r/K mixtures. a) Growth curves for different populations from the spatial computational model. The yellow line represents the r-cell population, the green line represents the mixture population of r- and K-cells and the red line represents the K-cell population. The Y-axis represents population size. The X-axis represents time. (n = 100 stochastic simulations per population; mean ± SD). **b) Mean growth rate comparison among populations.** The growth rate was measured at 1 hour intervals. The Y-axis represents mean growth rate. The X-axis represents time. (n = 100 stochastic simulations per population; mean ± SD, Student’s t-test: ****P<0.0001). **c) Necrotic area detection.** The second column represents the necrotic area in xenografts. **d) Proportion of the non-necrotic area (y-axis) in xenografts. e) Whole tumor (black) and viable cell (gray) weight in xenografts.** The xenografts were extracted at the sixth week after cell inoculation. n=6 for each xenograft type; Student’s t-test: *P<0.05, ****P<0.0001.

#### In vivo

Mouse xenografts initiated with r, K, and r-K mixed cells were weighed on the 34^th^ day, followed by H & E staining. The necrotic and non-necrotic regions were distinguished using the gray threshold method. We observed a high incidence of death in the r-xenografts (Figure 5c) and a significantly higher proportion of non-necrotic cells the mixed xenografts (Figure 5d). Although average fresh weight of the r-xenografts is much larger than the fresh weight of K- and mixed xenografts (reflecting the higher r cell proliferation rate, Figure 5e), the mean weight of viable cells in the mixed xenografts is the highest. It indicates that the existence of phenotypic trade-offs within a cell population is advantageous to cell viability and population growth.

## Discussion

*r*- and *K*- selection theory predicts that natural selection increases density-dependent rates of population growth. This has been tested experimentally on specific model systems from bacteria and *Drosophila* to vertebrates accounting for life history details^15,70–73^. The notion of trade-offs in life-history evolution became a prominent feature of the theory and prompted a focus of theoreticians and field scientists both in ecology and evolutionary biology^1,2,15,37,74^. However, the heart of continuing controversy on the theory of *r*- and *K*-selection between theoreticians and field biologists is that many complex life-history characters of natural populations contradict theoretical expectations^1,4,5^. It is unrealistic to expect that a theory could account for all aspects of the natural environment and its impact on evolutionary processes in all organisms^1,5^. Thus, the only proper way to test the theoretical predictions is in controlled settings congruent with the assumptions of the simple models.

Tumorigenesis is an evolving and dynamic process where highly genetically and phenotypically heterogeneous neoplastic cell populations persist in challenging environments. In fact hallmarks of cancer cannot be acquired in all cancer cells all the time^75^. An important cell-to-cell phenotypic variability is determined by several exterior and interior constrains^12,76–79^. For instance, environments in tumors are both stable (but crowded, hypoxic, and nutrient-poor) in the interior, and fluctuating in nutrients, space, and interaction between the components in the microenvironment at the edge of the tumors^80,81^. The consequences of somatic cell evolution under complex environmental pressures parallel ecological processes in nature, with inevitable survival-reproduction trade-offs because organisms have to allocate limited resources among several functions that affect fitness. Neoplastic cells may also be subject to evolutionary trade-offs with respect to resource allocation and growth constraints^12,20,21,35^. The mixture of biotypes that form cancer cell populations can be characterized by survival-proliferation trade-offs, and directly quantified in controlled environments *in vitro*. Carrying out experimental evolution under *r*- and *K*-selection in cancer cell lines, we observe that cancer cell populations face a survival-reproduction trade-off. r cells are selectively favored to allocate the majority of their resources to reproductive activities at the cost of their ability to propagate under crowded conditions, consistent with the central idea of the *r*- and *K*-selection theory^4^.

Our analysis of pathway enrichment and expression of differentially expressed genes reflects phenotypic differences in cell proliferation, cell death, and adhesion between r and K cells *in vitro* and *in vivo*. We observe higher growth and death rates in r cells, compared to K cells (Figure 1d and 1g, Figure 2b). Additionally, adhesion junctions and focal adhesion affect adherence capability of cells. Since trypsin is frequently applied to dissociate adhesive cells from their substratum^82^, we performed a trypsinization assay to quantify cell adhesive ability (Extended Data Figure 6). Extended Data Figure 14 shows that it takes significantly longer to digest attached K than r cells, confirming that K cells are more prone to adhesion.

Cells with higher fitness tend to maintain a relatively high transcriptome stability^83^. Changes in transcriptional profiles reveal that r cells are much more sensitive to density change than K cells, consistent with the observation that r cells have lower fitness at high density in competition assays (Figure 1b and 1c, Extended Data Figure 1a and 1b). Remarkably, differentially expressed genes that respond to density change in r cells are enriched in the cell cycle and DNA replication pathways (Extended Data Figure 7, 8), suggesting that r cells have different growth rates depending on culture conditions. This is consistent with direct measurements of growth rate at high and low density (Figure 1d).

Computer simulations which integrate of *r*- and *K*-selection theory predictions and parameters of inter-cell interaction based on Lotka–Volterra models illustrate temporal and spatial dynamics of population growth of heterogeneous cell populations following *r*- and *K*-strategies. The growth curves based on empirical data and mathematical models show that growth rates and fitness of r- and K-selected cells follow the logistic equations predicted by theory. As density increases, K cells dominate mixed cell populations. Our simulations, fitted to empirical data, establish a competitive relationship between phenotypically diverse cancer cells. In the short term, competition can decrease whole-population fitness. However it triggers niche differentiation leading cell types to occupy different niches, thus maximizing the use of available resources in the ecosystem^60^. Interaction between tumor cells further improves the total fitness of a tumor in the long term (Figure 5).

## Materials and Methods

### Cell line

The HeLa cell line was provided by the Cell Bank, Type Culture Collection Committee, Chinese Academy of Sciences. The test for mycoplasma contamination was negative. The HeLa-HPV18 single-nucleotide variants^84^ were identified in the cell line. The HeLa cells were cultured in complete DMEM (Gibco) medium containing 10% FBS (Gibco) and antibiotics (100 μg/mL streptomycin and 100 units/mL penicillin, Sigma-Aldrich) at 37 °C in an atmosphere of 5% CO_2_.

### Cell cryopreservation

Cells were first trypsinized using a 1X trypsin-EDTA solution at room temperature for three minutes and suspended in complete growth medium. Suspended cells were collected by centrifugation (1300 rpm, 5 min) and resuspended in 1X PBS. PBS suspended cells were collected by centrifugation (1300 rpm, 5 min) and resuspended in cryopreservation medium. The cryopreservation medium contains 10% DMSO and 90% FBS. Cryopreservation medium suspended cells were pipetted into a cryopreservation vial gently, and placed into a −80 °C freezer. Finally, vials were transferred intoliquid nitrogen for long-term storage when temperature decreased to −40°C.

### Subculture and Single-cell isolation

Cells were washed with 1X PBS three times after discarding cell culture medium, and trypsinized with 1X trypsin-EDTA solution at room temperature for three minutes. The detached cells were suspended, divided, and transferred into plates. Single cells were sorted into individual wells of 96-well plates by flow cytometry (BD) from a HeLa cell population. After six hours, a microscopic examination was performed to ensure only one cell in a well.

### eGFP and dsRed transfection

Cells were transfected by Lentiviral vectors pLenti6.3-MCS-IRES-eGFP and pLenti6.3-MCS-IRES-dsRed (Invitrogen). Approximately 5 × 10^6^ HeLa cells were incubated in a 10cm dish with DMEM before transfection. After incubating for 24 h, the DMEM medium was replaced by 10 mL transfection-mix-medium that contains 8 μg/mL polybrene and 10^6^ IU/mL lentivirus particles. The multiplicity of infection (MOI) value was 1. After transfecting for 24 hours, cells were washed with PBS three times. To select cells that stably express eGFP and DsRed, the transfected cells were cultured in DMEM medium with blasticidin (10 µg/mL) for at least four weeks.

### Density-dependent selection

#### Evolution experimental system

The initial cell population (IN-cells) derived from a single cell which was randomly selected from the HeLa cell line. When the population size of IN-cells reached 10^7^, it was randomly divided into two sub-populations of equal size. Each of sub-population was labeled with fluorescent proteins as described above. Density-dependent selection was performed on labeled cells.

#### r-selection

Cells were cultured under low-density. To ensure low density, cells were seeded on the surface of a 10 cm dish with approximately 128 *cells*/*cm*^2^. Every 120 hours when the population density reached to about 4 × 10^3^ *cells*/*cm*^2^, a subset of cells was transferred to a new plate to keep a similar density as the original population (128 *cells*/*cm*^2^). Six replicates (three with dsRed and three with eGFP) were maintained in this manner for almost 200 cell generations (200 days). Samples from each population were cryopreserved in liquid nitrogen every 40 days.

#### K-selection

Cells were cultured under high-density. To ensure high density, cells were seeded on the surface of a 10 cm dish with approximately 10^5^ *cells*/*cm*^2^. Every 72 hours, when the population density reached to about 2.2 × 10^5^ *cells*/*cm*^2^, a subset of cells was transferred to a new plate to keep a similar density as the original population (10^5^ *cells*/*cm*^2^). Six replicates (three with dsRed and three with eGFP) were maintained in this manner for almost 130 cell generations (200 days). Samples from each population were cryopreserved in liquid nitrogen every 40 generations.

### Relative fitness assay

To measure the relative fitness of two cell populations cultured under a specific cell density (routine-, r-, or K-), the two cell populations were mixed and cultured together. The proportions of the two populations were monitored by flow cytometry (BD, Ex/Em (nm): 346/442) once a subculture was performed. Time between two subcultures depended on the culture protocol. The higher fitness population is the one dominating the mixed population over time.

### Measurement of growth rate

Population growth rate was estimated using the equation (1):

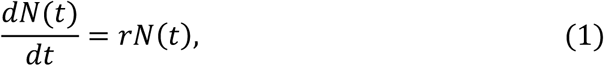

where *dN*(*t*) is the total number of cells at time *t*, and *r* is a constant coefficient. To obtain a linear function, cell numbers were converted to base-2 logarithms. The least-squares method (LSM) was used to fit the linear regression with the slope (r) of the regression line estimating the growth rate.

### Soft agar colony formation assay

Approximately 1,000 cells were suspended in a top layer of 0.4% soft agar (SeaPlaque Agarose, BMA products). The cell suspensions were then overlaid onto a bottom layer of 1% soft agar containing complete DMEM supplemented with 10% FBS in six-well plates. After a week, colony number was counted. After three weeks, the images of colonies were collected to compare their diameters by microscopy.

### *In vivo* tumor growth

Female BALB/c Nude mice were purchased from the Beijing Vital River Laboratory Animal Technology Co., Ltd. Mice were 5–10 weeks of age for all experiments and kept in germ-free environments in the Institute of Zoology, Chinese Academy of Science.

Five-week female BALB/c Nude mice were assigned randomly into cages upon arrival. IN-cells, r-cells, and K-cells were suspended in normal saline separately. 50 μL cell suspension (2 × 10^5^*Cells*/*μL*) was inoculated under the inguinal skin of the mice. For each type of cell inoculation, three mice were randomly selected and sacrificed every week from the third week after inoculation. Xenografts were collected for further analysis. Sample sizes were determined empirically (based on experience of other investigators who did similar assays). The experiments were not performed blind. All animal study protocols were reviewed and approved by the review boards of the Institute of Zoology Animal Care and Use Committee, Chinese Academy of Science (ethical approval reference number IOZ-20150061) and were conducted in accordance with the National Institutes of Health Guidelines for the Care and Use of Laboratory Animals. The maximal tumor diameter of 20 mm was permitted by Institute of Zoology Animal Care and Use Committee, Chinese Academy of Science. None of the experiments in this study exceeded this limit.

### FACS analysis of G0/G1 phase and cell death/apoptosis

Cells were trypsinized and suspended in cold 1X PBS.Five μL propidium iodide (Sigma, P4170) was added to the suspension. Cells were incubated at 4 °C for 30 minutes. Stained cells were analyzed by flow cytometry (BD, Ex/Em (nm): 346/442). Data were collected from 10000 stained cells.

The Annexin V, Alexa Fluor® 350 conjugate (Invitrogen, a23202) was used for apoptosis rate detection. Cells were trypsinized and diluted to ~1 × 106 cells/mL in the annexin binding buffer. Five μL annexin V conjugate was added to 100 μL of the cell suspension. The cell suspension was incubated at room temperature for 15 minutes. After the incubation, 400 µL annexin-binding buffer was added. The samples were kept on ice after mixing gently. The stained cells were analyzed by flow cytometry (BD, Ex/Em (nm): 346/442). Data were collected from 10000 stained cells.

### Necrotic area detection and calculation of the weight of viable cells in mouse xenografts

The H&E (haematoxylin and eosin) staining of tumor sections was used to detect necrotic cells^85^. First, we prepared the central H&E staining section of a xenograft. The sections were then converted into digital images using Aperio Digital Pathology Slide Scanner (Leica). To detect necrotic areas in a xenograft, the images of sections were read via Matlab (MathWorks) and converted to gray scale (rgb2gray function in Matlab). Image contrast was enhanced using histogram equalization (histeq function in Matlab). We then adjusted image intensity values twice with parameters low_in=0.1 and high_in=0.7 (imadjust function in Matlab) and applied the 2-D median filtering to the image with the filtering parameters m=5 and n=5 (medfilt2 function in Matlab). Finally, we set grey scale value 90 as the threshold to distinguish the necrotic and non-necrotic areas of the image. The pixels in the tumor region with the grey scale value less than 90 were considered necrotic. The net weight of viable cells in a xenograft tumor was obtained by multiplying the total weight of a tumor by the proportion of the non-necrotic area.

### RNA-seq and data analysis

Total RNA was isolated using the TRIzol reagent, as described by the manufacturer (15596018, Invitrogen). RNA-seq libraries were constructed and sequenced by Berry Genomics. RNA-seq NGS reads were aligned to the hg19 reference genome using the Mapsplice aligner (version 2.1.8)^86^ with default parameters. The gene-level expression levels were quantified by RSEM (Version 1.2.19)^41^, based on the TCGA mRNA-seq Pipeline (https://webshare.bioinf.unc.edu/public/mRNAseq_TCGA/UNC_mRNAseq_summary.pdf). Differentially expressed genes between samples were detected using EBSeq (Version 1.1.5)^87^ and were defined as the PPDE over 0.99. Gene set enrichment analysis of KEGG pathways was performed using the Functional Annotation Tool from DAVID with default parameters^88,89^. Expression perturbations in significant KEGG pathways were determined by GAGE ^46^ with default parameters.

### Trypsinization assay

Cells were seeded on a six-well plate. After 12 hours, we discarded the culture medium and washed the cells with cold PBS three times. 500µL 0.05% Trypsin was added into the well at room temperature. The plate was swayed softly and slowly 20 times. All supernatant (about 500µL) was transferred into a new tube. We pipetted the supernatant gently to make sure most cells were single individuals. 400µL of the supernatant was put back on the plate to trypsinize the remaining cells. Finally, we estimated cell numbers in the 100 µL of the remaining supernatant (*N*_1_) and among the remaining cells (*N*_2_) using a hemocytometer. The following equation was used to calculate the trypsinised cell ratio within a time interval:

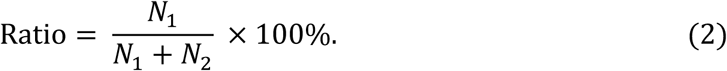

### Immunofluorescence assay

Cells were seeded on coverslips. The coverslips were then placed on six-well plates. After a 48-hour incubation, cells were fixed with 4% paraformaldehyde (PFA) in PBS for 20 minutes at room temperature, followed by blocking and permeabilizatuion for 30 minutes in blocking buffer, comprising 2% bovine serum albumin (BSA) and 0.2% Triton X-100 in PBS. Cells were incubated with the Yap1 antibody (GTX35195, GeneTex) for one hour, then with the FITC-conjugated goat anti-rabbit IgG (H+L) polyclonal antibody (GTX 77059, GeneTex) for 30 minutes. Both antibodies were diluted in PBS with 2% BSA. Cell nuclei were stained with Hoechst 33342 (H3570, Invitrogen). Images were acquired using a Leica TCS SP8 confocal laser microscopy system (Leica Microsystems).

### Real-time quantitative PCR with reverse transcription

Total RNA was isolated using the TRIzol reagent, as described by the manufacturer (15596018, Invitrogen). 1 μg of RNA was used to generate cDNA with the High Capacity cDNA Reverse Transcription Kit (4368814, Applied Biosystems). Real-time quantitative PCR was performed to amplify cDNA by using Maxima STBR Green/ROX qPCR Master (K0223, Thermo Scientific) in a CFX96 Touch Real-Time PCR Detection System (Bio-Rad). The average threshold cycle (Ct) of quadruplicate reactions was determined and amplification was analyzed by the ΔΔCt method. Gene expression was normalized to that of GAPDH. Real-time quantitative PCR with reverse transcription data were representative of at least three independent experiments, with two technical replicates per experiment. Primer sequences used to amplify human DLG2 and GAPDH as follows:

human DLG2 forward: CAATGGGATGGCAGACTTTT;

human DLG2 reverse: ACAGCTCGGTGGAGAAACAT;

human GAPDH forward: ACAGCCTCAAGATCATCAGC;

human GAPDH reverse: ATGGACTGTGGTCATGAGTC.

### siRNA knockdown

siRNAs (Lipofectamine 3000 transfection reagent) were used to knock down the expression of DLG2. To check the knockdown efficiency, total RNA was isolated and quantified by quantitative PCR (qPCR) three days after transfection. The target sequences used to knock down human DLG2 are as follows:

si-h-DLG2_001: ACCUCAUUCUUUCCUAUGA;

si-h-DLG2_002: GCUAGAACAAGAAUUUGGA;

si-h-DLG2_003: GGAGAUGAAUAAGCGUCUA.

### r/K competition assay *in vitro*

K cells with dsRed and r cells with eGFP were mixed together equally. Cell density of the mixture population was about 2×10^6^ cells/mL. 500 μL of cell mixture was loaded on the central surface of an empty culture plate. After five minutes, the plate was put back to the incubator. When all cells attached (almost two hours), sufficient complete growth medium was added to the plate. Microscopic fluorescent field images of the plate were collected using imageXpress XLS (Molecular Devices) every three days. Image data were analyzed following the pipeline in the imageXpress XLS data analysis software (Molecular Devices).

### Model fitting

Density dependent population dynamics can be predicted using a variety of mathematical models^90,91^. The logistic and Gompertz growth models are most frequently used^92,93^. To determine which mathematical model is suitable for us to predict cell population dynamics, we first obtained cell population dynamics data over eight days via the MTT cell proliferation assay, then fit population dynamics curves to three models: Logistic, Gompertiz, and Exponential (Extended data Figure 9). We created a nonlinear model for cell population growth based on the data from the MTT assay (fitnlm function in Matlab). The adjusted *r*-squared value of the logistic growth curve is 0.856, the Gompertz – 0.828, and exponential – 0.739. This suggests that the logistic growth model fits the data best.

### Density-dependent population growth model

We chose the Lotka-Volterra (L-V) model to investigate population dynamics cell type mixtures^64,94^. We assume that the cell population 1 and cell population 2 are two sub-types of cells from the same population. These two types of cells compete for the same resources in a mixted population. The competitive Lotka–Volterra equations are

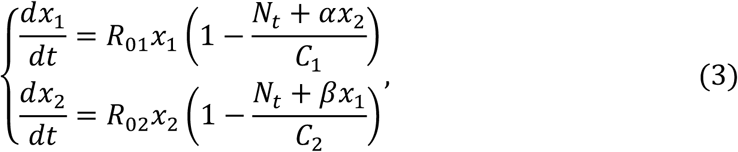

where *x*_*i*_ is the size of the ith population; *R*_0*i*_ is the inherent per-capita growth rate, and *C*_*i*_ is the carrying capacity. *α* represents that the additional effect of cell population 2 on cell population 1 and *β* represents the additional effect cell population 1 on cell population 2. *N*_*t*_ represents the total cell number at time *t*. Note, the meanings of *α* and *β* are slightly different from the competitive coefficients in the general Lotka–Volterra equations. *α* + 1 and *β* + 1 are equivalent to competitive coefficients in the general Lotka–Volterra equations.

Per-capita growth rate (**R**_**0**_) estimation: The inherent per-capita growth rate of every cell must be known at the beginning as the population growth model calculates the growth rate of every cell and its progenitors separately. It is an easy way to evaluate the inherent per-capita growth rate of every cell at the beginning via random sampling if the inherent per-capita growth rate distribution of a cell population is known. We isolated 141 single cells from the K-cell population and 100 single cells from the r-cell population and cultured them separately in wells of 96-well plates. Cells were counted every day for each clone over five days. The growth each cell over five days is considered exponential because cell density is very low.

Growth rate was estimated using the equation 4:

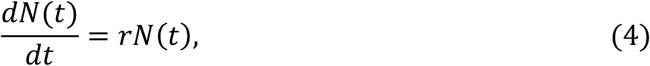

where *dN*(*t*) is the total number of cells at time *t*, and *r* is a constant. To obtain a linear function, cell numbers were converted to base-2 logarithms. The least-squares method (LSM) was used to fit the linear regression in which the slope (r) of the regression line estimates the growth rate.

We assume that mutations do not drastically affect the growth rate immediately. Therefore, r is equivalent to the R_0_ of the initial single cell. We assume that the growth rate of any given type of cell comes from a specified normal distribution. We estimate distribution parameters from empirical growth rates of 141 K cells (for K cell simulations) and from 111 r cells. (Extended Data Table 5).

The distribution of the inherent per-capita growth rate of a cell population is *R*_0_~Norm(*μ, σ*^2^) and lies within the interval *R*_0_ ∈ (−∞, +∞). The parameters of the inherent per-capita growth rate distributions are estimated using the function ‘*normfit*’ in MATLAB. The fitted distributions are *DR*_0*r*_~Norm(1.1832, 0.2441^2^) and *DR*_0*K*_~Norm(0.6823, 0.3764^2^).

Carrying capacity (C) estimation: Given the logistic cell population growth curve, carrying capacity can be estimated using the logistic growth function. We seeded r and K cells separately on six-well plates separately and assessed population size every 24 hours. The initial population size was 1.5 × 10^3^cells (Extended Data Figure 10).

Since apoptotic cells could not be distinguished when the cell counting was performed each day, cell number may have been over-estimated. Severity of over estimation of r-cell number grows as cell density increases because these cells go into apoptosis at a high rater as conditions become crowded. It is necessary to correct the estimation of carrying capacity to eliminate the effect of apoptotic cells on cell count. The carrying capacity could be corrected by

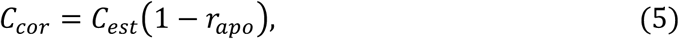

where the C_Cor_ represents the corrected carrying capacity, C_est_ represents the estimated carrying capacity via curve fitting, r_apo_ represents the r- or K-cell apoptosis rate under high density (data were collected via FACS analysis of cell apoptosis).

The carrying capacity of r cells was estimated as *C*_*r*_ = 1.937 × 10^4^ *cells*/*cm*^2^ and of K cells as *C*_*K*_ = 2.2216 × 10^4^ *cells*/*cm*^2^.

**α** and **β** estimation: α and β directly influence population size. Cell growth rates can decrease rapidly α and β are both large, leading to population sizes far beyond carrying capacity. However, empirical observations did not show significant reduction of population size when *r-* and *K*-cell were mixed together at high density. It suggests that either α or β is very small. The r- and K cells co-cultural assays suggest that K cells may have a higher competitive ability (Figure 4). This inference indicates that β should be near 0. Here we assume that β is between 0 and 0.5 and α between 0.5 and 3.

First, we use a grid-search scheme, all parameter pairs traversed with intervals 0.5, to estimate the α and β. We predict the dynamics of an *r* and *K*-cell mixture population using the density-dependent growth model with set values of α and β. Other parameters were fixed. We then calculated Pearson correlation coefficients between predictions and observations. These correlations are maximized when (α, β) = (2.5,0). This fit is better than when (α, β) = (2,0) or (α, β) = (3,0) (Extended Data Table 4). The correlation values increase and then decrease within the interval α ∈ [2,3] when β = 0. This suggests that the values of α between 2 to 3 and β = 0 maximize the agreement between simulations and data.

We next repeated the grid-search scheme, traversing values of α between 2 and 3 with intervals of 0.1 and setting β = 0. The final estimates are: α = 2.2 and β = 0 (Extended Data Figure 5;11, Extended Data Table 4).

Cell population dynamics of the mixed population with among-cell competition are based on the values of *C*_*r*_, *C*_*K*_, *C*^*r*^/*α* and *C*^*K*^/*β*. WE estimate C_r_ = 1.937 × 10^4^ *cells*/*cm*^2^, C_K_ = 2.2216 × 10^4^ *cells*/*cm*^2^, α = 2.2, and *β* = 0. Thus, *C*^*r*^/*α* = 9.685× 10^3^ and *C*^*K*^/*β* is infinite. Here *C*_*K*_ > *C*^*r*^/*α* and *C*_*r*_ < *C*^*K*^/*β* indicate that r cells would eventually go extinct when competing with K cells^95^.

### Spatial growth model

Tumor cells living in a limited space cannot move freely. Among-cell interactions are also confined to a limited space, precluding interaction when between-cell distance is large. Given these considerations, we assume that there is a cell-centric limited space for every cell where the density-dependent effects which impact the central cell are confined. We call this density-dependent space (DDS; for more details see Stochastic simulation of cell growth with spatial structure).

When a population grows logistically, its growth is exponential early on, provided it carrying capacity is much greater than its size. However, if carrying capacity is small, early-stage population size increase results in a drastic decrease in growth rate. Carrying capacity is related to the size of the habitat. In a DDS, the maximum carrying capacity is 36 (Extended Data Figure 13, methods: Carrying capacity estimation of spatial growth model). Here we use *C*_*s*_ to represent the carrying capacity in a DDS. Because the *C*_*s*_ value is very small (compare to *C* in equations (3)), the equations (1) are not applicable to predict the dynamics of r- and K-cell mixed population using the spatial growth model. In addition, only when cell density exceeds a certain threshold, do the cells become subject to density-dependent growth. A reasonable value of the threshold is 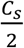 as the population growth rate achieves its maximum when the population size reaches 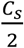.

For two cell sub-populations, we let

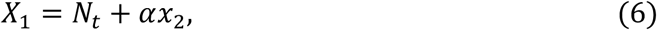

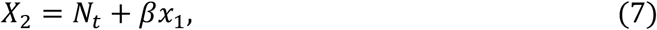

where *X*_1_ represents the practical population size which determines the density-dependent growth rate of population 1; *X*_2_ is defined similarly for population 2.

When 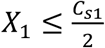, the growth rate of population 1 is equal to its inherent growth rate, and similarly for population 2 when 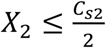. For population 1, we have

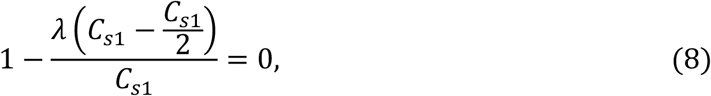

Where the *λ* is a constant. By solving the equations 7, we get *λ* = 2. The final equations are

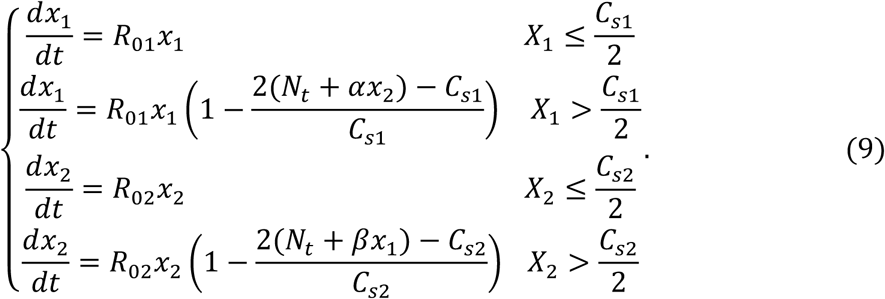

Carrying capacity estimation of the spatial growth model: In the spatial model, we assume that the density-dependent space (DDS) is a square area which contains 36 grid coordinates (6X6 grids, Extended Date Figure 10). Because the carrying capacity of *K*-cells is 1.147 times that of *r*-cells (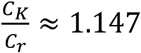; more details see the carrying capacity estimation), in a DDS the carrying capacity of *K*-cells is *C*_*sK*_ = 36 (the maximum number for the region), and of *r*-cells is *C*_*sr*_ = 31.

### Stochastic simulation of population growth of r and K cells in co-cultures

Cells in culture are subject to artifacts, such as subculture. A subculture is performed when cell density exceeds a threshold (roughly 70% to 90% confluent) and is used to maintain cell density. The subculturing procedure includes recommended split-ratios and cultural medium replenishment schedules. A realistic *in silico* cell culture model should take into account such artifacts. The details of the stochastic simulation procedures are as follows:

#### Initiation

We assign the initial inherent per-capita growth rate to every cell on initialization. Here we assumed that growth rates of cells in a population come from a normal distribution. Every cell is assigned an initial growth rate sampled from its growth rate distribution (see the per-capita growth rate estimation for details). To avoid outliers, random sampling was based on a truncated distribution (within the interval *R*_0_ ∈ (Q1, Q366 of the inherent per-capita growth rate. Q1 is the lower quartile and Q3 the upper quartile of the observed growth rate distribution respectively (Extended Data Table 5). The initial population size was chosen according to culturing methods being simulated and followed experimental conditions.

#### Population growth and sub-culture

Cell division is based on growth rate. Each cell in a population, enters the division stage only if the cell growth rate is over 0. In the stochastic simulation, the time of a cell cycle (CT) was defined as

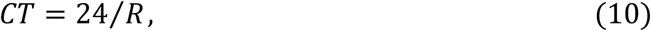

where the *R* represents the growth rate of a cell calculated from the density-dependent population growth model. CT is measured in hours. The biological meaning of *R* is the number of cell divisions within 24 hours. Considering the characteristics of the cell cycle, the time of interphase of mitosis occupies nearly 90% of the entire cycle^96^. Thus, cell division time (DT) is:

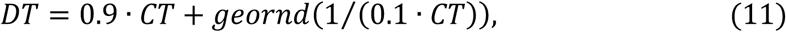

where *geornd* means a random number sampled from a geometric distribution. If the time since the last cell division is greater than DT, the cell divides into two cells. When the culture time exceeds 72 hours (*K*-selection conditions), subculture was performed. Population size was reduced to the initial population size during subculture by random selection. After subculture, cells continue to grow *in silico*.

Six mixed populations with different r- and K-cell initial proportions (r:K=99:1, r:K =9:1, r:K =7:3, r:K =1:1, r:K =3:7, r:K =1:9) were simulated. 100 simulations were performed for each population type (Extended Data Figure 12).

### Stochastic simulation of population growth with spatial structure

Tumor cells live in a spacially heterogeneous environment^97,98^. The distribution of cell density within a tumor should therefore be non-uniform. Spatial structure plays an important role in population dynamics^31,61,99–105^. In a given cell growth space, if the real-time location of cells can be determined, the spatial structure of the whole population can be described. For this reason, we constructed a two-dimensional lattice-based growth space where physiological activities such as cell growth and migration are carried out. The location of cells is determined by grid coordinates. (Extended Data Figure 13)

To simulate the population dynamics of cells which grow on a two-dimensional surface as realistically as possible, we considered the following factors that can influence spatial structure: cellular morphological characteristics, cell migration, cell proliferation, and cell death.

#### In-silico cellular morphology

The growth of cells on a two-dimensional surface may result in regional differences in cell density due to uneven cell distribution or different growth rates (Extended Data Figure 14a). In other words, the density-dependent spatial heterogeneity exists in the cell growth environment. In addition, space occupied by cells varies under different densities. In a low-density environment, cells occupy a larger area, increasing cell surface to maximize contact with the culture medium. Because of crowding, cells are arranged closely. The attachment area of a cell decreases in a high-density environment (Extended Data Figure 14b). Therefore, we assume cells have two in-silico morphological types: large cell, corresponding to cells growing in a low density environment, occupying four coordinate grid; small cell, corresponding to cells growing in high density environment, occupying one coordinate grid (Extended Data Figure 13). The *in silico* cellular morphology can be transformed between large and small cells. When there is an empty coordinate around a small cell that can accommodate a large cell, the small cell preferentially transforms into a large cell. A large cell will switch its morphology to two small cells via cell division when and only when there is no space around it that can accommodate two large cells, and there is space to accommodate two small cells.

#### Cell migration

Cells migrate with a certain rate in their growth space. When a migration event occurs, the coordinates of cells in the growth space change once, and the migration must occur to adjacent coordinates (Extended Data Figure 13). The migration direction of each step is assumed random. Cells differ in their migration speed. r cells migrate more readily than K cells, as measured by a trans-well migration assay. The mean migration speed of r-cells is close to five times higher than K cells (Extended Data Figure 15). Here we assume that the migration speed follows a beta distribution. The parameter “a” of the beta distribution is 5. The expected migration speed of r cells is 0.5 and of K cells is 0.1.

#### Cell proliferation

A division event can only be completed in two adjacent coordinates. Cells retain their original cellular morphology during division. If there is no space to proliferate, small cells die.

#### Cell death

When a cell dies, its original coordinate is marked as empty and can be occupied by another cell via cell division or migration. Death occurs if a cell that must divide but has no space to do so, or if a cell is affected by high density (calculated by Equations (9)).

#### Density-dependent space

Density-dependent space (DDS) is a square region containing 36 grids (6X6 grids). A cell can be in the center (large cells) or on the grid coordinate (3,3) whose origin is the top-left of the DDS grid (small cells). We assume that only the cells located in the DDS contribute the density effects to the central cell (Extended Data Figure 13).

#### Simulation process

The simulation program of cell growth with spatial structure is divided into two processes: initiation and population growth. In the initiation process, cells were loaded in the center of the growth space. All cells were clustered together to form a circle community. This constructs a density-dependent spatially heterogeneous environment for cell growth. The outside space low-density, while inside the cell community is a relatively high-density environment. The inherent per-capita growth rate (Normal distribution; the same as the initiation process in the stochastic simulation of cell growth with subculture) and migration speed (Beta distribution) of each cell were initiated with random sampling. Finally, the program calculates a constant variable δt, which represents the minimal time interval that can contain a migration event. In the population growth process, the migration and proliferation of cells depend on the migration rate and the density growth rate (calculated by Equations (9)). The migration rate and the inherent per-capita growth rate were maintained between mother and daughter cells. The program iterates all cells and calculates their density growth rates. δt is the time interval between iterations.

### MTT assay

Cells were suspended and seeded at the concentration of 500 cells/well in 96-well plate. A volume of 20 μl dissolved MTT was pipetted into each well. After incubating for 4 h at 37 °C in a humidified CO_2_ incubator, the medium was removed and 200 μl sterile DMSO was added to each well. Absorbance values were then read at 570 nm with a microplate spectrophotometer. The proliferation of living cells was monitored based on absorbance values.

### Migration assay

Migration assay was performed using 6.5mm Transwell inserts (3422, Corning) containing polycarbonate membrane filters (8-μm pore size) for 24-well plates. Briefly, cells were digested with 0.05% trypsin, and suspended in a FBS-free DMEM culture medium. Cells were then plated into the upper chamber (3×103 cells/well). At the same time, 650µL of DMEM with 10% FBS was added to the lower chamber of the well and the plates were incubated for five hours at 37 °C with 5% CO_2_. After incubation, cells on the upper surface of the membranes were removed gently with cotton swabs. Cells that had entered the lower surface of the filter membrane were stained with 0.1% Hoechst 33342 (H3570, Invitrogen) for 30 minutes at room temperature, and washed three times with PBS. Four randomly selected fields in each well were image captured with the ImageXpress Micro HCS (Molecular Devices), and migrated cells were counted. n = 3 independent experiments.

### Statistical analyses

Statistical analyses were performed using R. Student’s *t*-test and Wilcoxon test were used for calculating significance of between-group differences. Statistical significance is indicated by P < 0.05. All data were expressed as mean ± s.d. of at least three independent experiments.

---

**Extended Data Figure 1.**
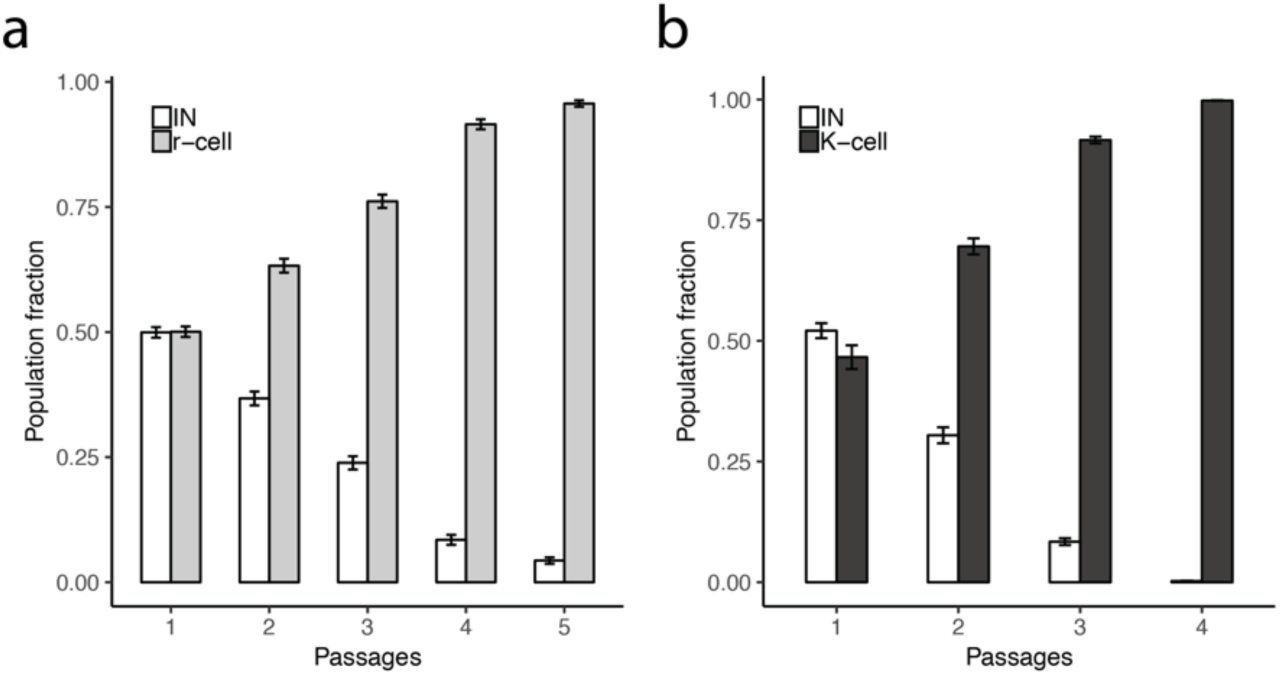
Fitness, growth, and apoptosis rates of r- and K-cells in 2D and 3D environments. IN cells and the a) r-cells or b) K-cells were mixed together at equal amounts at the beginning. The mixed populations were cultured under normal culture conditions. The proportion of each type of cells was measured by flow cytometry every two days during subculture. Error bars represent standard deviations. N= 3 independent experiments per population, mean ± SD.

**Extended Data Figure 2.**
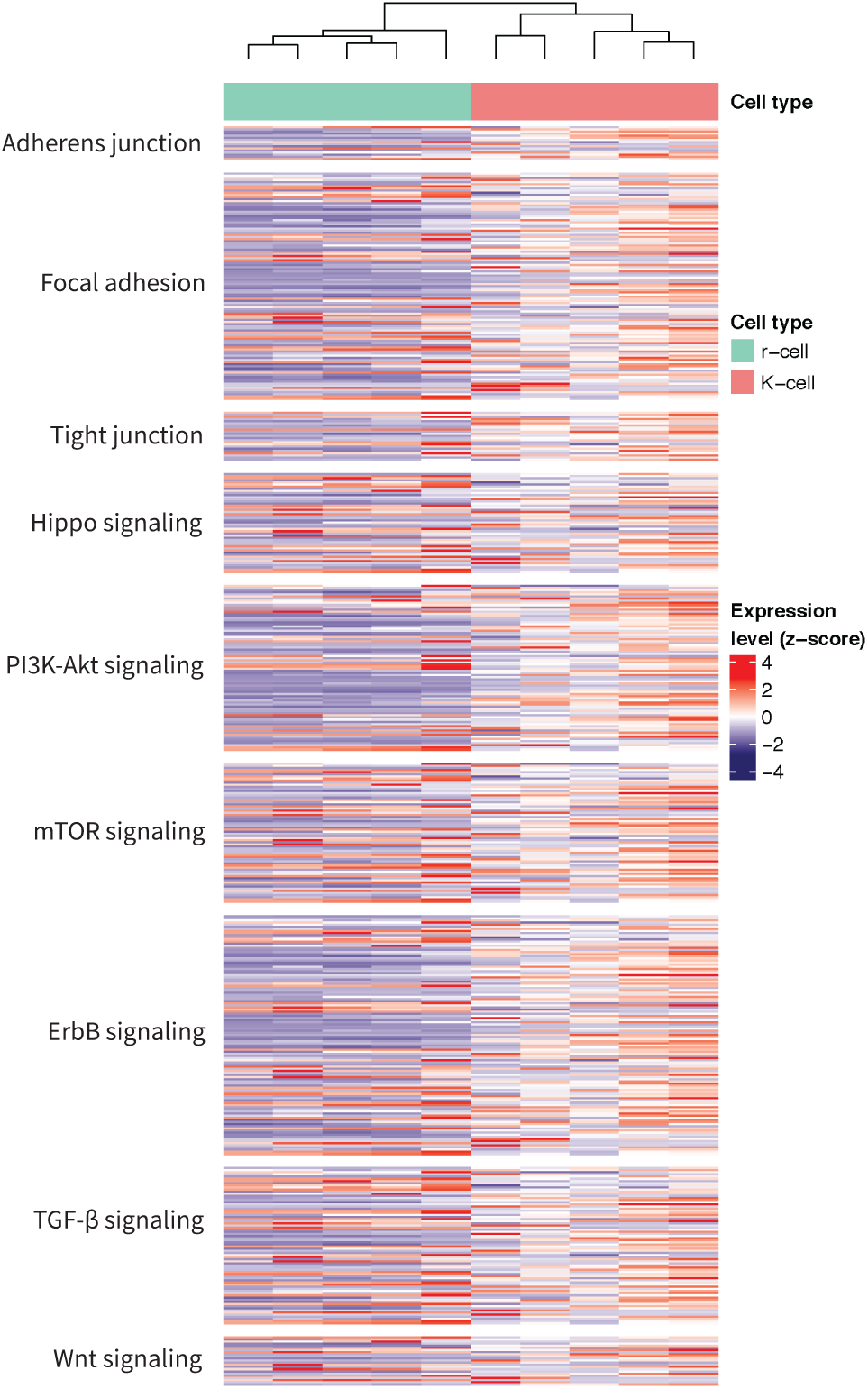
Cluster of Hippo-related pathways.

**Extended Data Figure 3.**
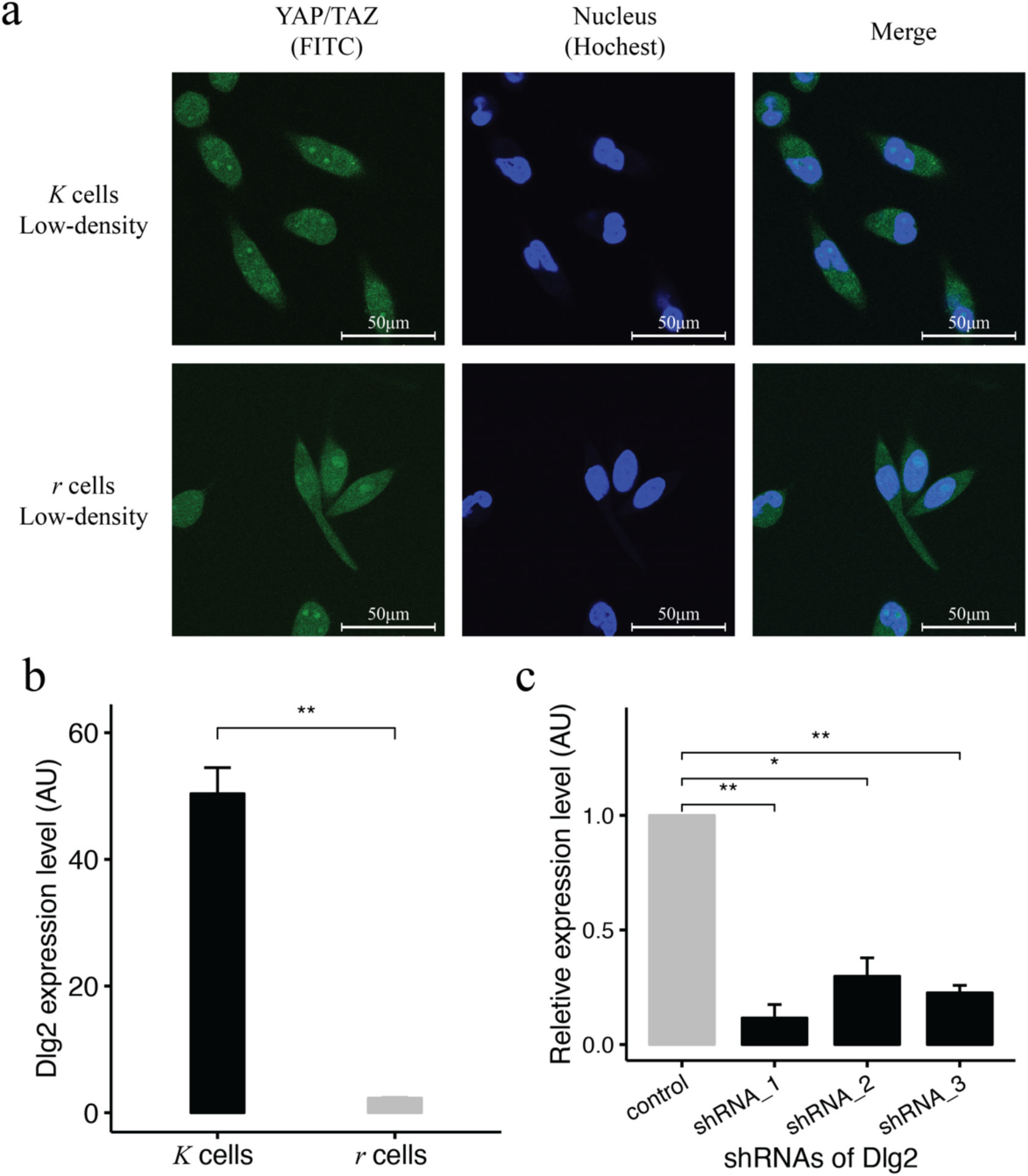
YAP/TAZ colocalization in r and K cells under low density and shRNA knockout of Dlg-2 in K cells. a) YAP/TAZ colocalization in the cytoplasm and nuclei at low density. YAP/TAZ were stained with FITC by immunofluorescence. Positive Hoechst staining marks nuclei. Scale bars represent 50μm. b) The expression level of Dlg-2 in r and K cells at high density by q-PCR. c) Dlg-2 shRNA in K cells. Three shRNAs were used for the Dlg-2 knockdown. n=3 independent experiments per population, mean ± SD.

**Extended data Figure 4.**
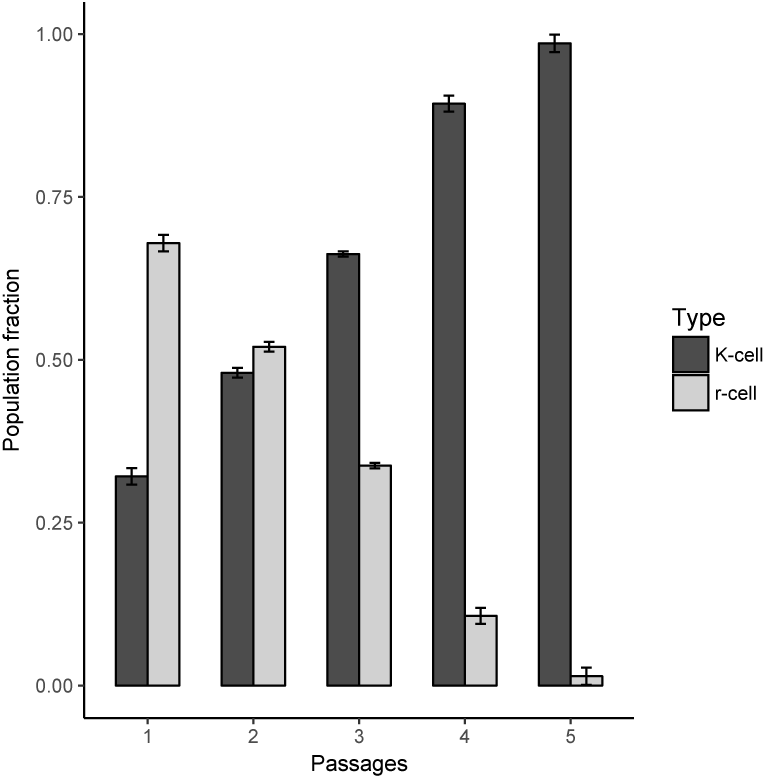
Observed dynamics of mixted populations initiated with 90% r-cells and 10% K-cells. Mixed populations were cultured under K-selection. The proportion of each type of cells was measured by flowcytometry every three days during. The bars represent the proportion change of cell population by time. The grey bars represent r- and black bars K-cells. The x-axis represents subculture times, the y-axis represents cell-type proportions. Three replicates were performed on each assay. Error bars represent standard deviations. n=3 independent experiments per population, mean ± SD.

**Extended Data Figure 5.**
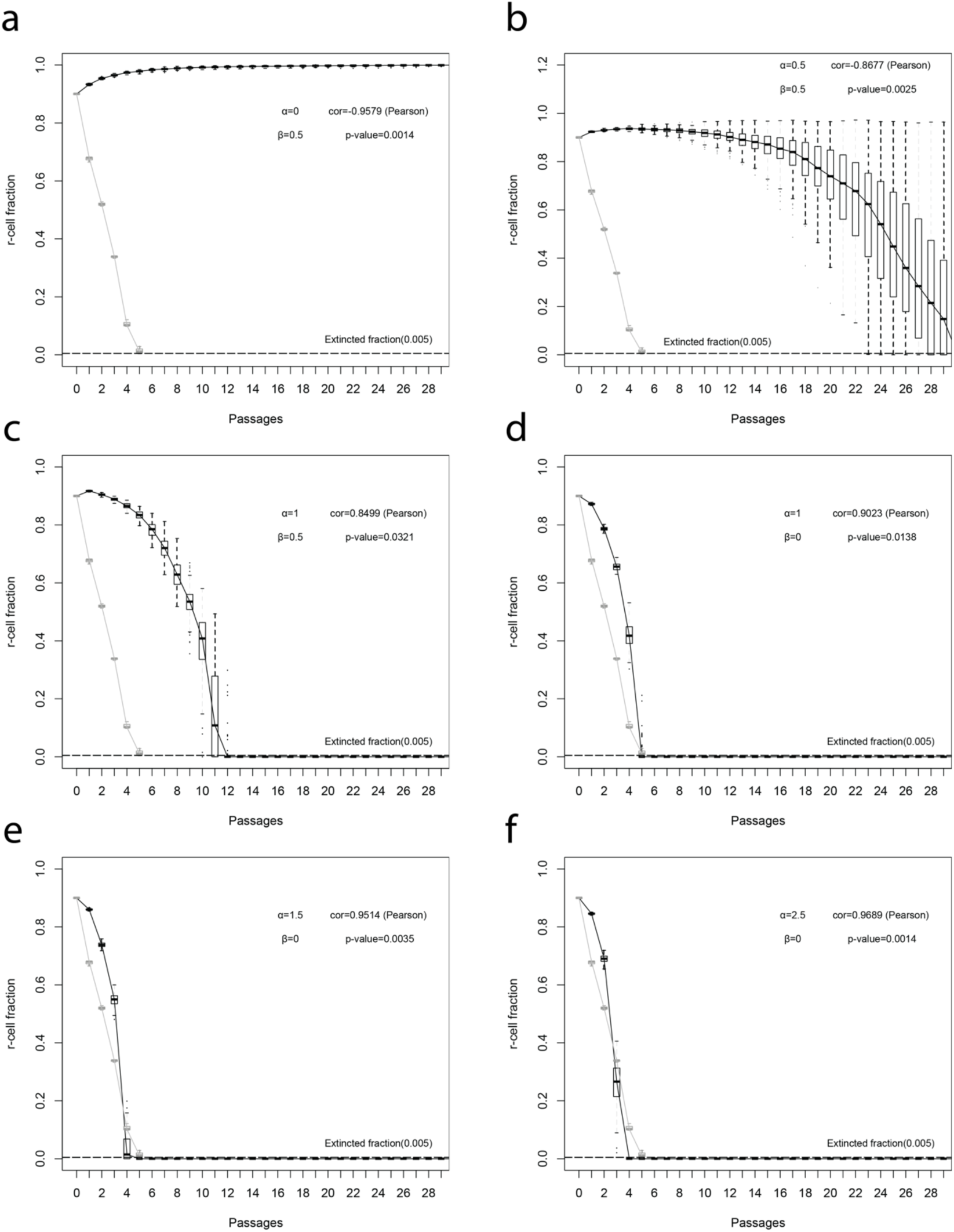
Predicted dynamics of r- and K-cell mixed populations. The proportion of each type of cells in the population was measured when subculturing. Sub-figures show the predicted population dynamics with different α and β. Black boxes and lines represent simulation results. Gray boxed and lines represent observations. n = 100 stochastic simulations per population, n= 3 independent experiments per population, mean ± SD.

**Extended Data Figure 6.**
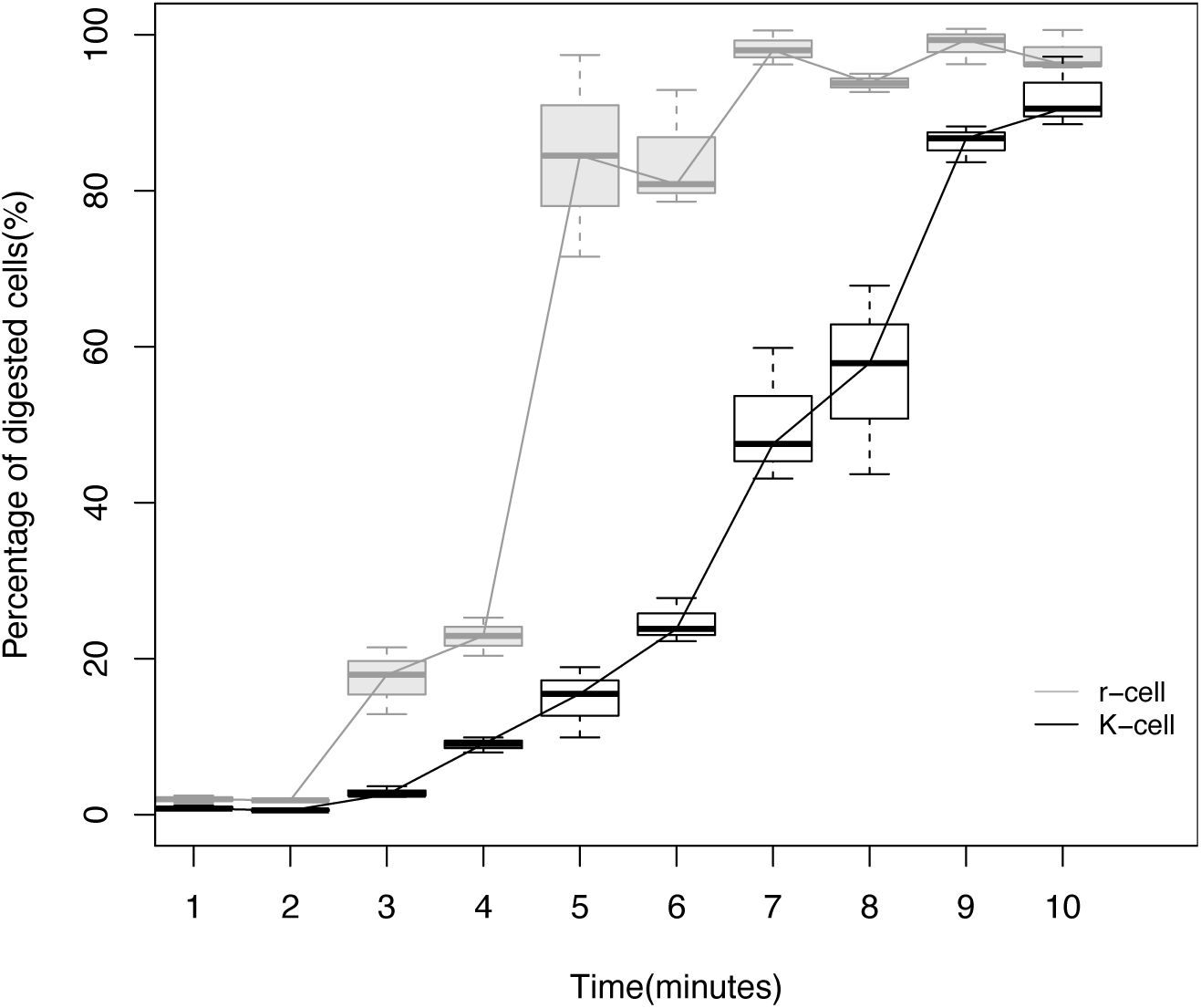
Detachment curves of r- and K-cells under trypsinization. Cells were digested by 1X Trypsin under room temperature. Cells which detached under trypsinization were counted every minute. The x-axis represents time and the y-axis represents the proportion of total cells that have been digested. Grey lines and box diagrams represent observations of r-cell populations. Black lines and box diagrams represent observations of K-cell populations. n= 6 independent experiments per population, mean ± SD. This figure shows that it takes significantly longer to digest attached K than r cells.

**Extended Data Figure 7.**
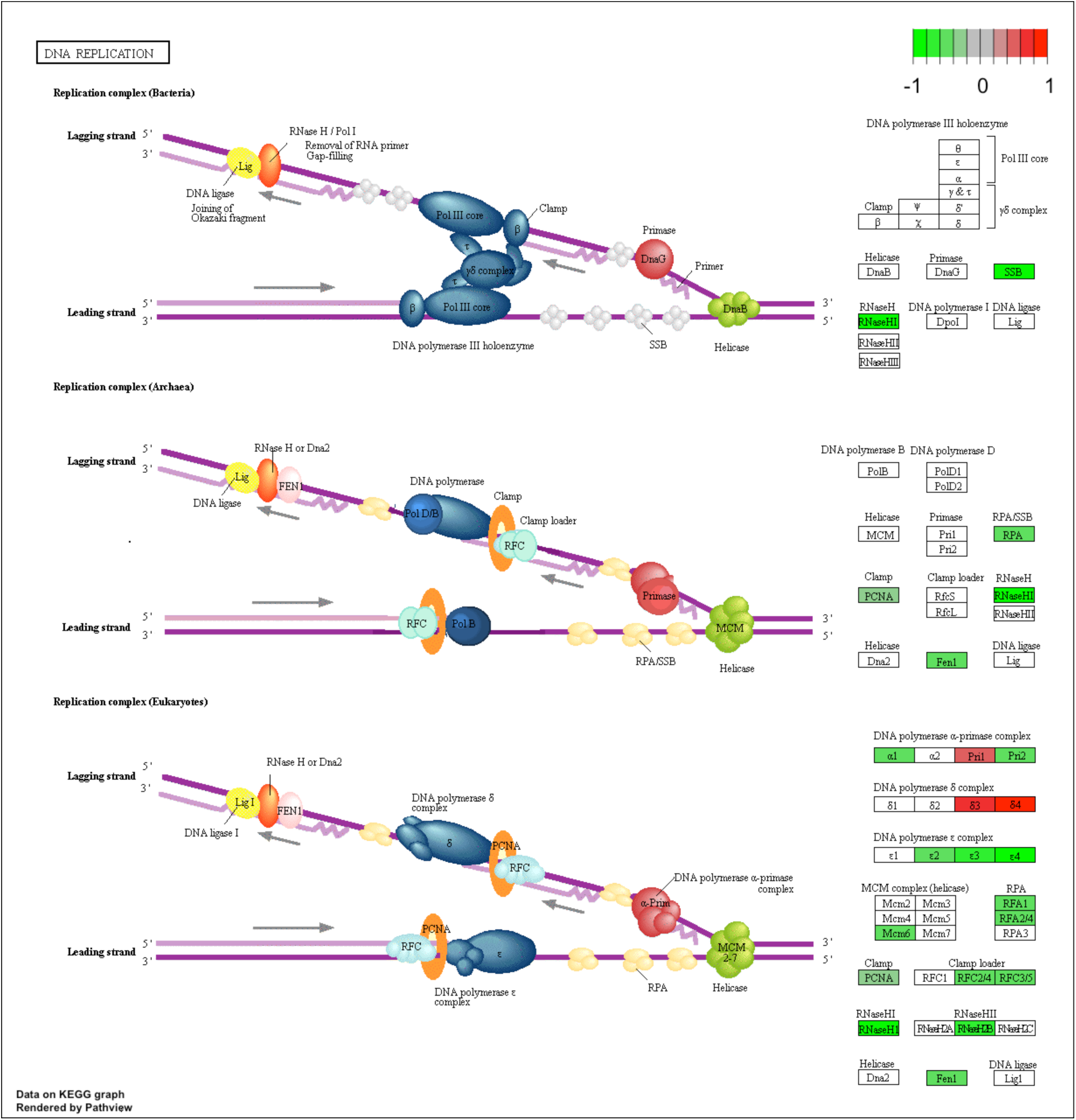
The DNA replication pathway is significantly lower expressed in r cells under high than under low density.

**Extended Data Figure 8.**
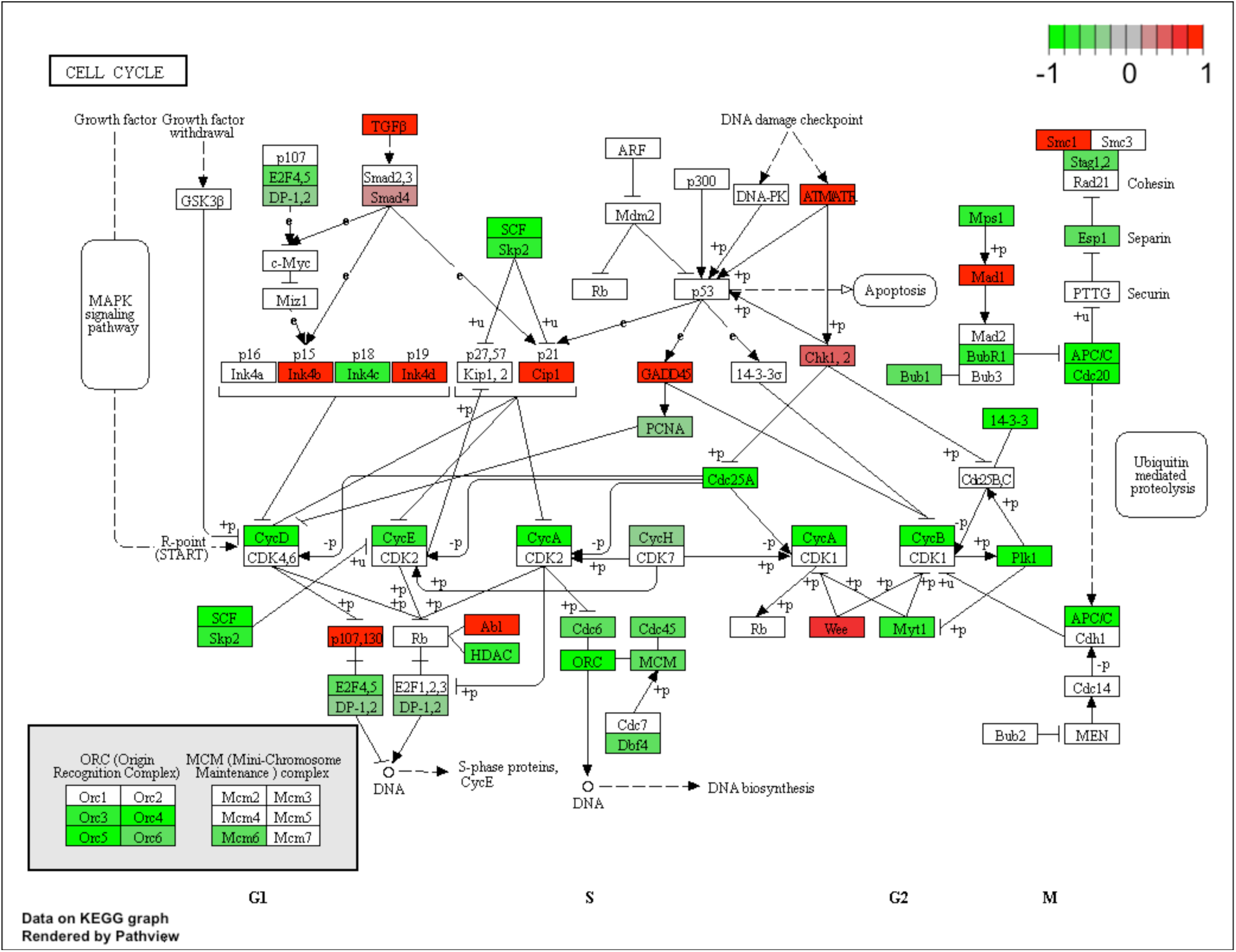
The cell cycle pathway is significantly lower expressed in r cells under high than under low density.

**Extended data Figure 9.**
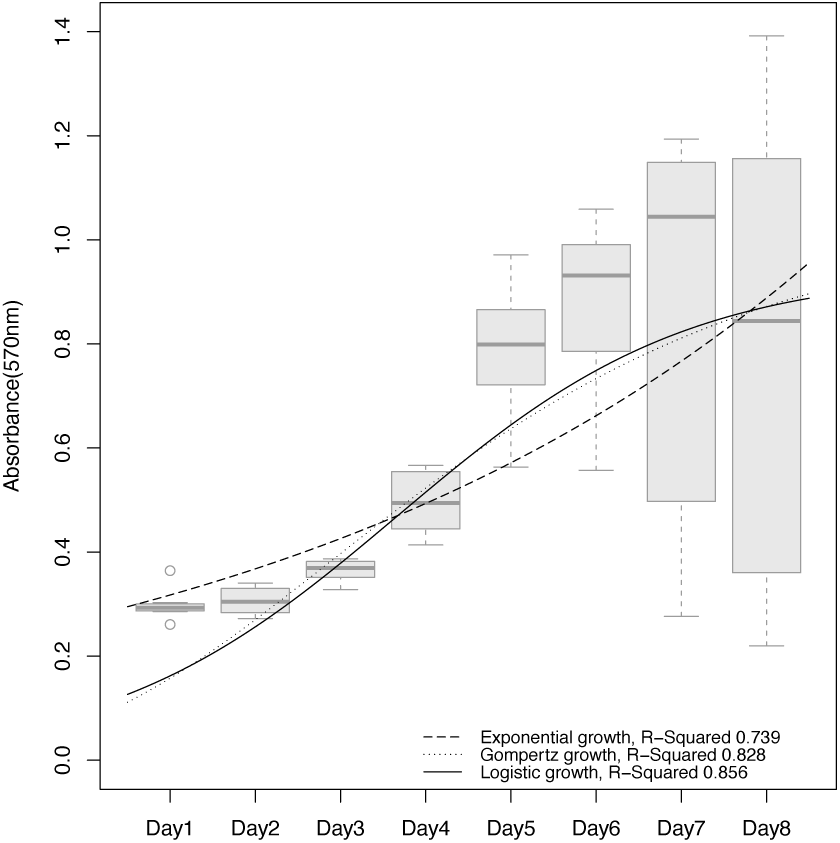
Growth model fitting. Cell growth was calculated using the MTT cell proliferation assay. We take the absorbance at 570 nm as the relative cell number. The assay was performed over eight days. We chose three population growth models: exponential, Gompertz, and logistic. Curves represent different models’ predictions.

**Extended Data Figure 10.**
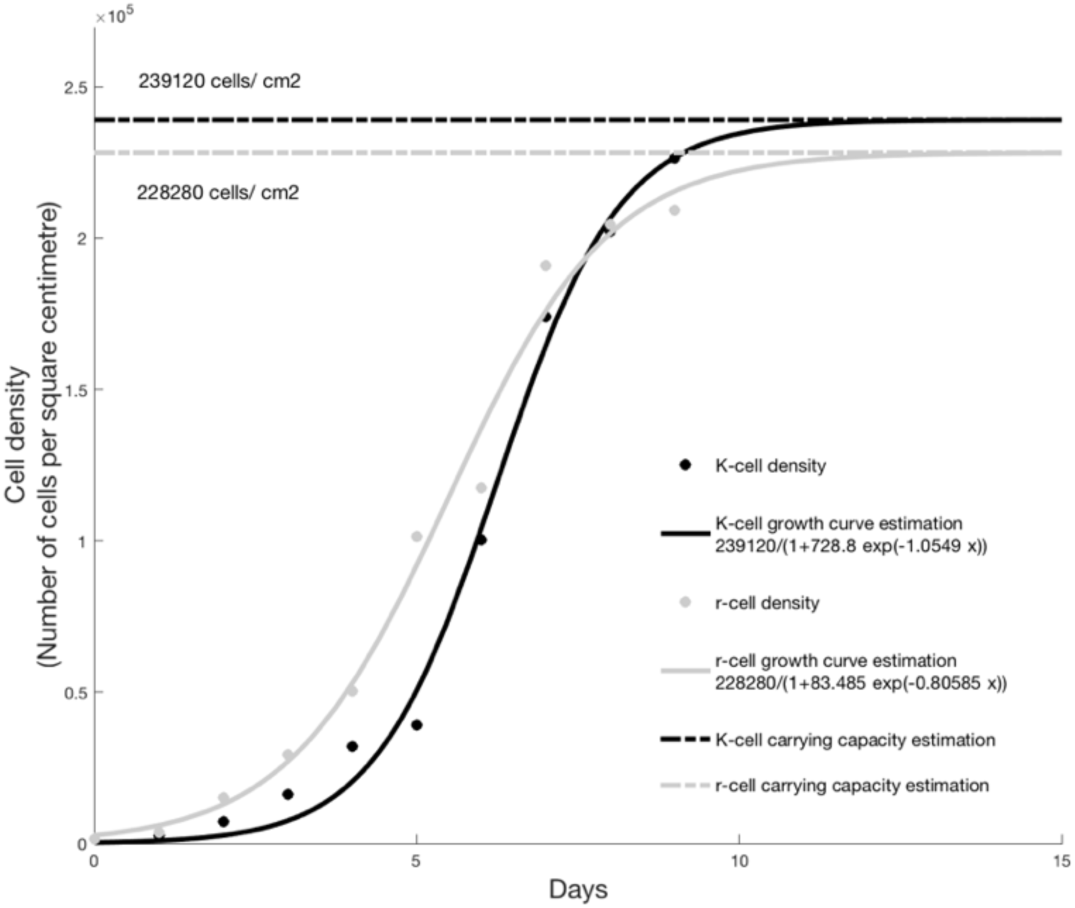
Carrying capacity estimation. The X-axis represents days after cell seeding. The Y-axis represents cell density. The unit of cell density is the number of cells per square centimeter. Grey points represent the cell density of r- and black points of K cells. Data were collected from experiments. Solid grey and solid black lines represent estimated growth curves of r- and K-cell populations respectively. Curve fitting was based on a logistic growth function. Parameters are shown in the legend. Adjusted R^2^ of the r-cell growth curve estimation is 0.985, for K-cells it is 0.991. The P-value of the r-cell growth curve is 2.21 × 10^−8^and for K-cell it is 6.4 × 10^−9^. The estimate of the carrying capacity is 239120 cells/cm^2^ for r- and 228280 cells/cm^2^for K cells.

**Extended Data Figure 11.**
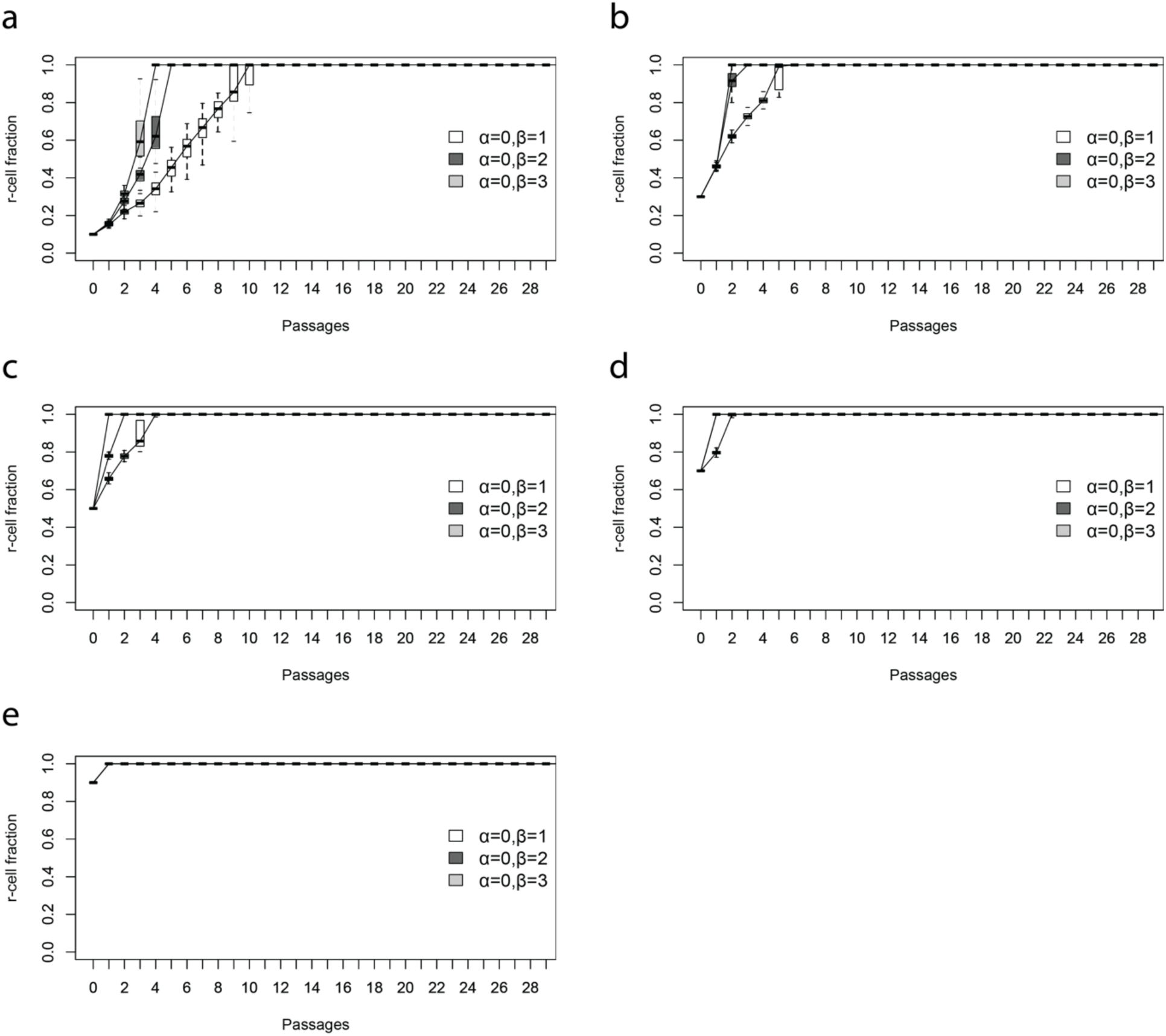
Predicted dynamics of r- and K-cell mixted populations. The mixed populations were cultured under high density based on the density dependent population growth model. The populations were initialed with cell density of 4 × 10^4^cells/cm^2^ and subcultured every 72 hours. The proportion of r cells is a) 10%, b) 30%, c) 50%, d) 70% e) 90% at the beginning. The proportion of each type of cells in a population was measured when subculturing. n = 100 stochastic simulations per population; mean ± SD.

**Extended Data Figure 12.**
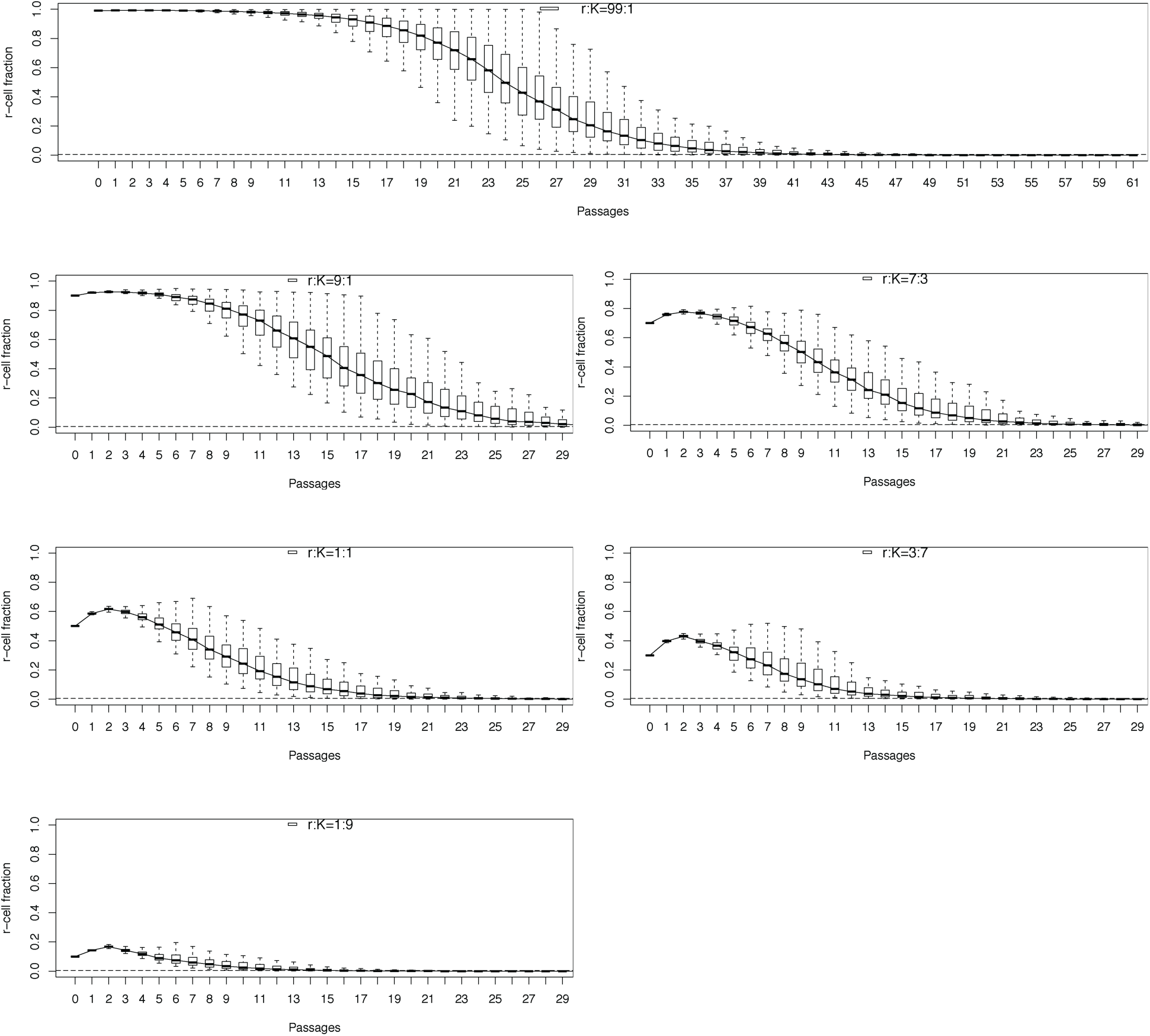
The dynamics of r- and K-cell mixture populations. Each panel shows 100 simulation predictions of a mixture population with a certain initial r and K cells ratio. The x-axis represents the passage times and the y-axis represents the r-cell fraction.

**Extended Data Figure 13.**
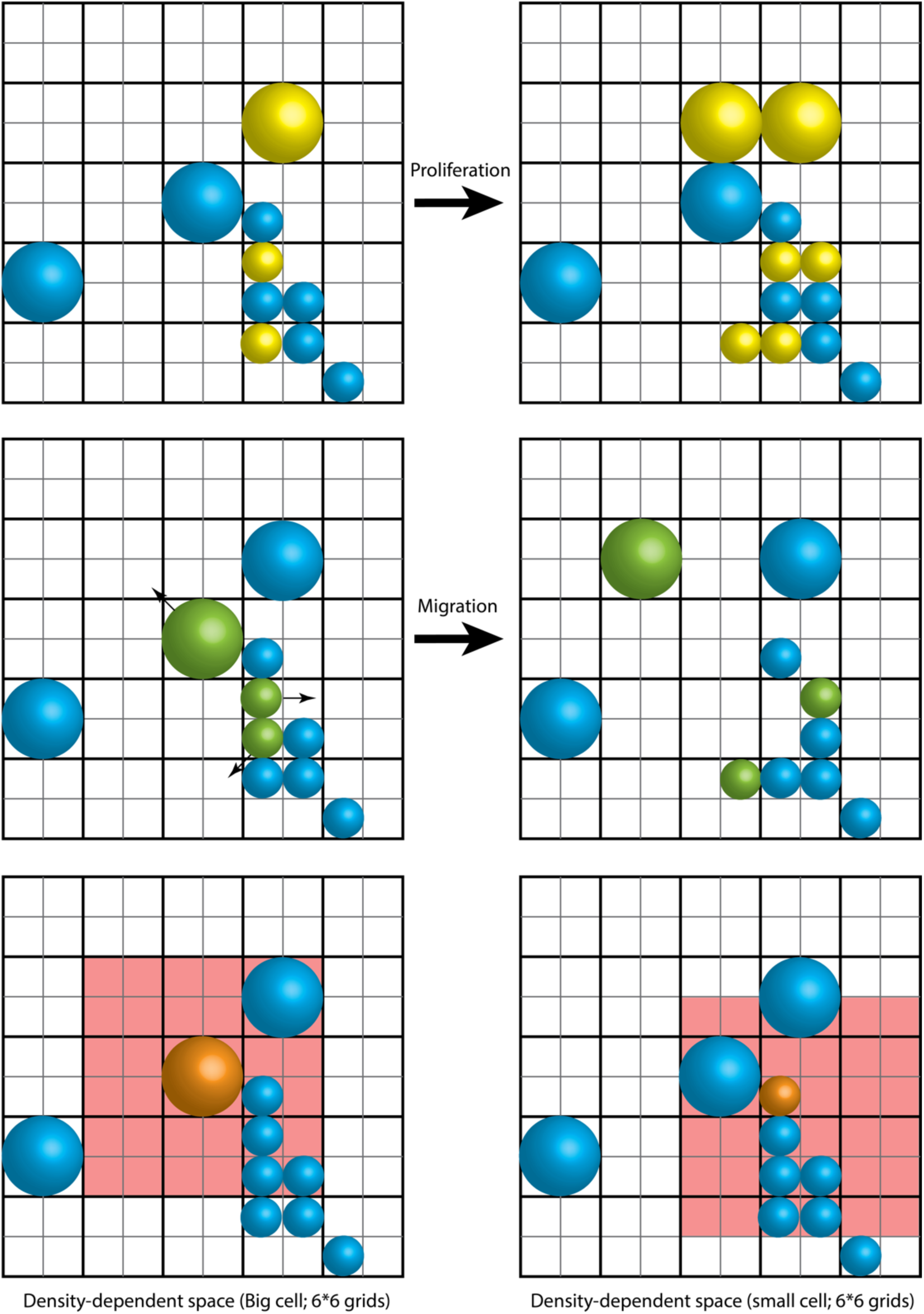
The spatial computational model of population growth. The cell growth space was assumed to be a two-dimensional planar grid. The location of cells is determined by grid coordinates. Cell migration and division are on the two-dimensional grid plane. The first line represents the division process. Yellow cells are undergoing mitosis. The second row represents migration with migrated cells in green. The red regions in the third row represent density dependent regions of migrated cells (orange).

**Extended Data Figure 14.**
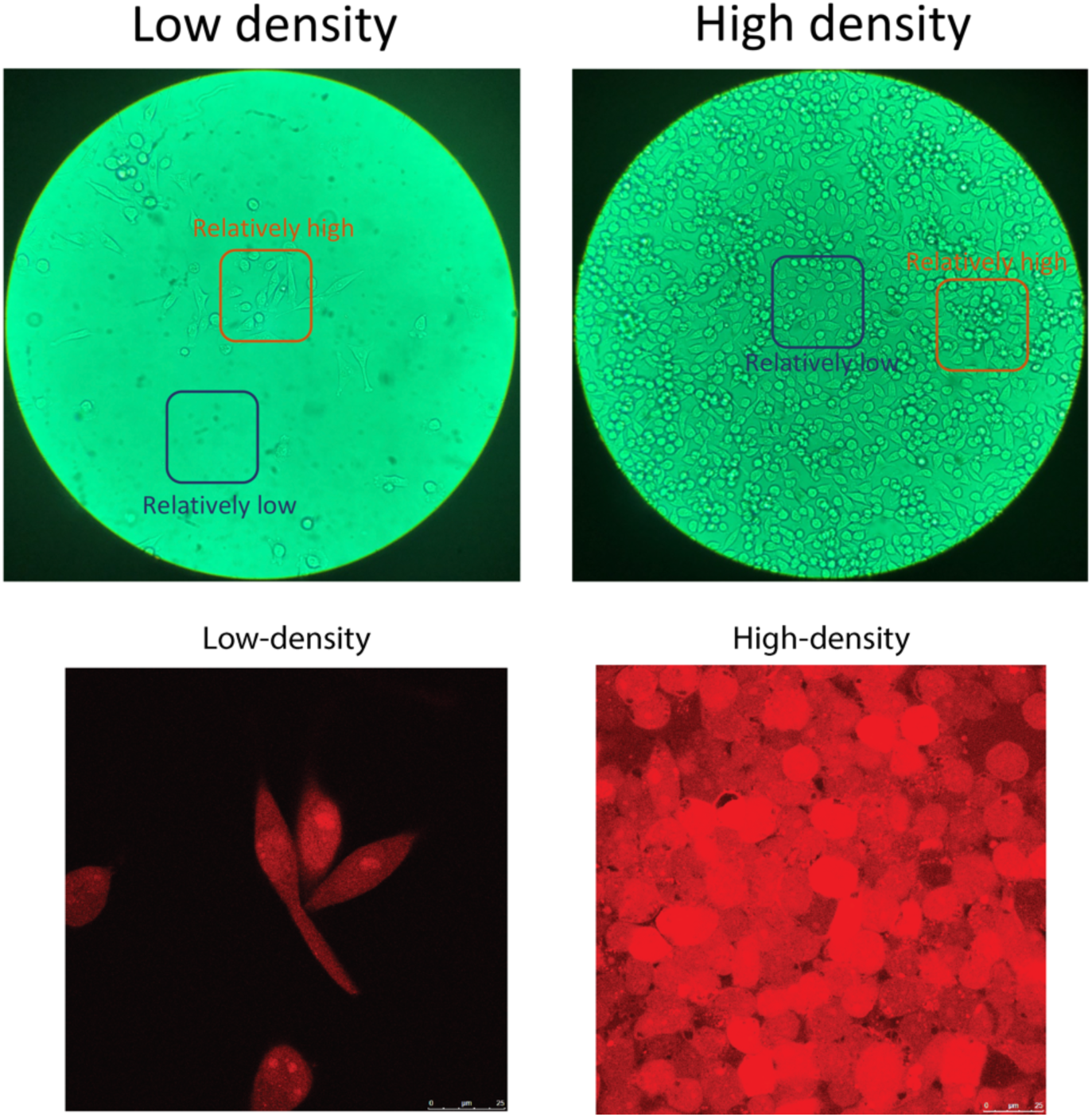
Density-dependent spatial heterogeneity and cell size. Upper: Images show density-dependent spatial heterogeneity across culture densities: low density on the left and high density on the right. Bottom: Fluorescence imaging of cells at two densities. Red marks cell bodies. On the left is the image of cells growing under low density and on the right under high.

**Extended Data Figure 15.**
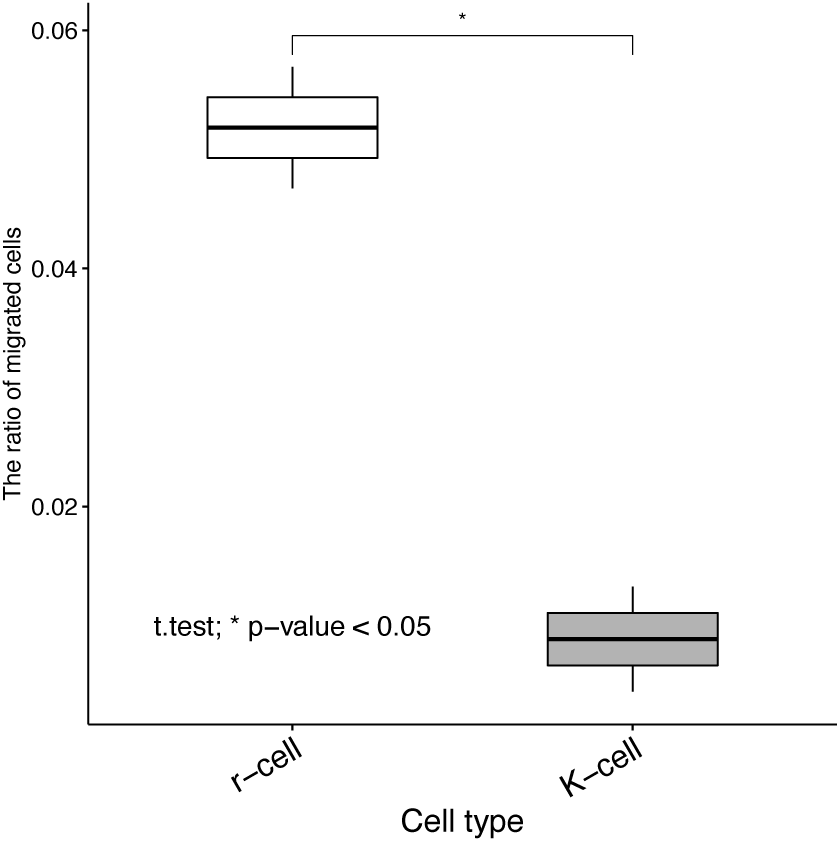
Ratio of migrated cells. r cells migrate more readily than K cells (t-test). The data were collected using a trans-well migration assay. n = 6 independent experiments; mean ± SD.

**Extended Data Table 1.** The number of DEGs across comparisons.

| Comparisons | High-expressed<br>genes number | Low-expressed<br>genes number | Total DEGs<br>number |
| --- | --- | --- | --- |
| KL vs. rL | 1748 | 1413 | 3161 |
| KH vs. KL | 1151 | 1126 | 2277 |
| rH vs. rL | 3284 | 3098 | 6382 |

**Extended Data Table 2.** Enrichment of DEGs in r- and K-cells under low-density.

| KEGG Pathway | Count | % | P-Value |
| --- | --- | --- | --- |
| Spliceosome | 54 | 1.7 | 1.00E-10 |
| Pathways in cancer | 97 | 3.1 | 2.90E-05 |
| Ribosome biogenesis in eukaryotes | 30 | 1 | 9.70E-05 |
| Small cell lung cancer | 29 | 0.9 | 1.60E-04 |
| Hepatitis B | 40 | 1.3 | 1.10E-03 |
| PI3K-Akt signaling pathway | 79 | 2.5 | 2.00E-03 |
| ECM-receptor interaction | 26 | 0.8 | 3.20E-03 |
| RNA transport | 43 | 1.4 | 5.20E-03 |
| Proteasome | 15 | 0.5 | 9.50E-03 |
| Base excision repair | 12 | 0.4 | 1.40E-02 |
| p53 signaling pathway | 19 | 0.6 | 2.30E-02 |
| Amoebiasis | 27 | 0.9 | 2.30E-02 |
| Epstein-Barr virus infection | 43 | 1.4 | 2.80E-02 |
| Pyrimidine metabolism | 26 | 0.8 | 3.20E-02 |
| Hippo signaling pathway | 35 | 1.1 | 3.50E-02 |
| Axon guidance | 30 | 1 | 4.20E-02 |
| Cell cycle | 29 | 0.9 | 5.10E-02 |
| Arginine and proline metabolism | 14 | 0.4 | 6.10E-02 |
| Amyotrophic lateral sclerosis (ALS) | 14 | 0.4 | 6.10E-02 |
| Influenza A | 38 | 1.2 | 6.30E-02 |
| Pertussis | 19 | 0.6 | 6.30E-02 |
| AMPK signaling pathway | 28 | 0.9 | 6.70E-02 |
| Purine metabolism | 38 | 1.2 | 7.20E-02 |
| RNA degradation | 19 | 0.6 | 7.80E-02 |
| Complement and coagulation cascades | 17 | 0.5 | 9.90E-02 |

**Extended Data Table 3.**
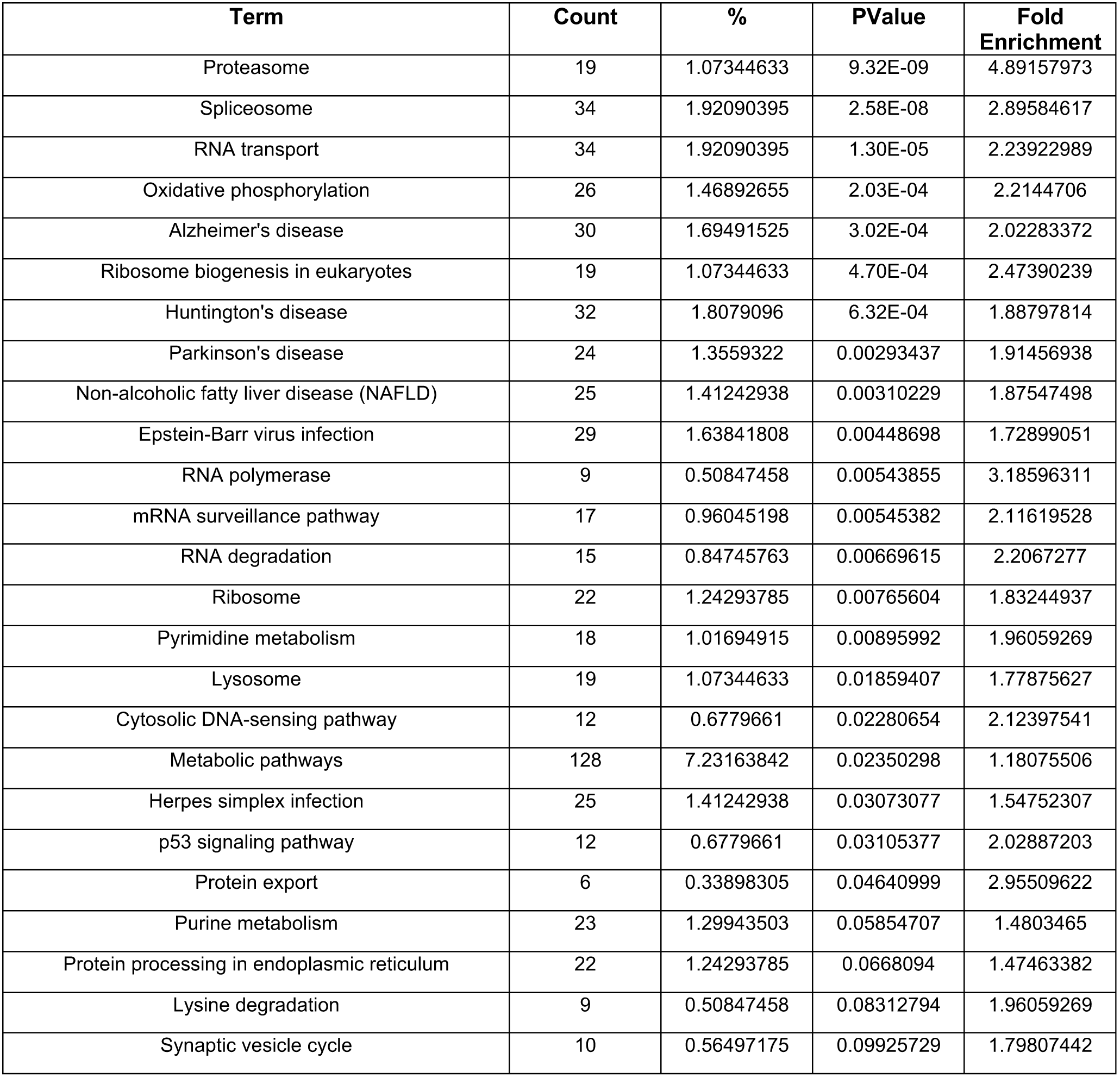
Top 25 pathways enriched in r- and K-cells under crowed culture.

**Extended Data Table 4.**
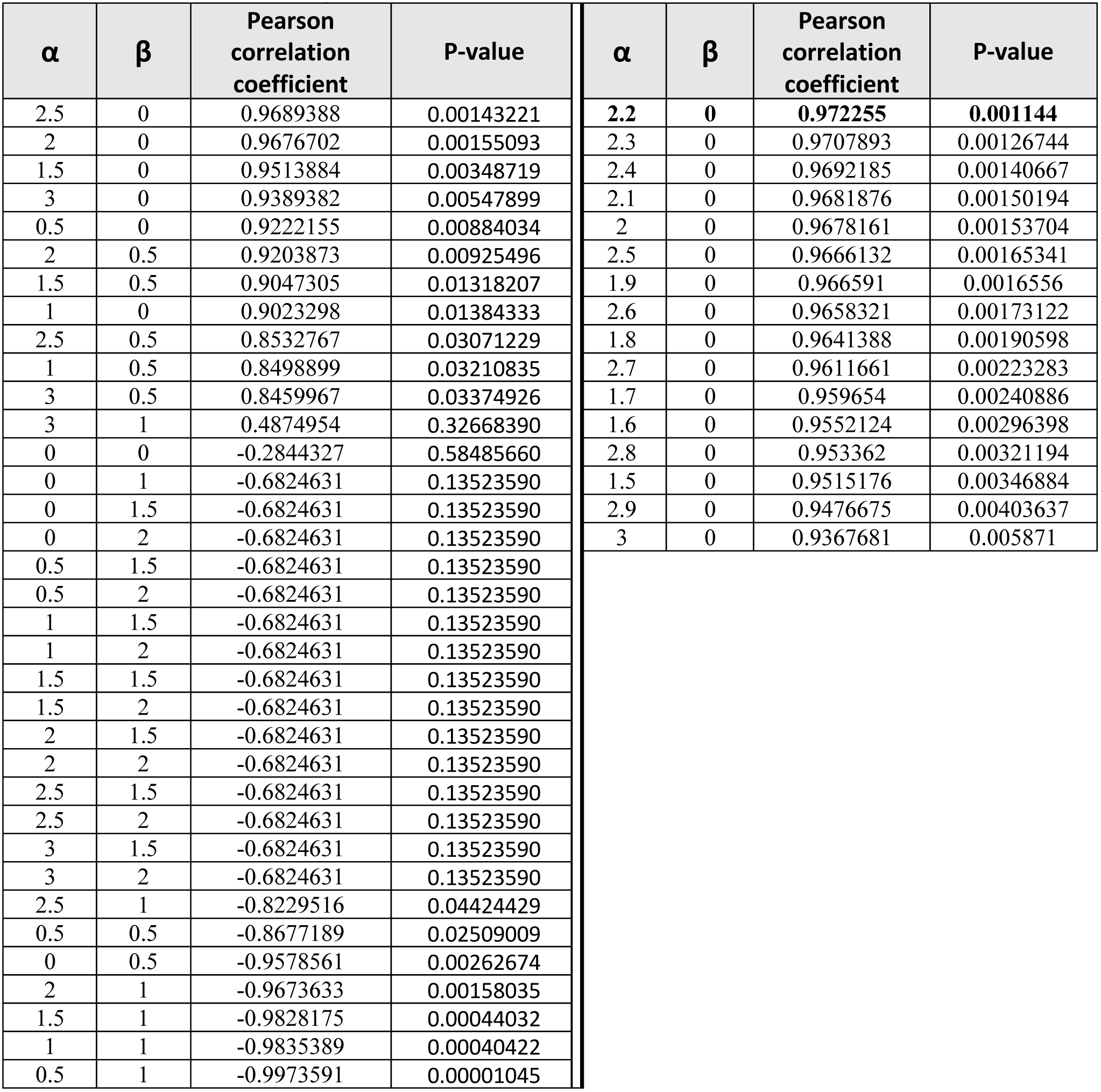
α and β estimation.

**Extended Data Table 5.** Single cell growth rate.

| Single cell growth rate |  |  |  |  |  |  |
| --- | --- | --- | --- | --- | --- | --- |
| Samples | IN G | IN R | H1G3KS | H1R1KS | H1G3RS | H1R1RS |
| 1 | 1.18205451 | 0.9099664 | 1.05785739 | 0.4169925 | 1.1529676 | 1.36096405 |
| 2 | 0.78450625 | 1.09276245 | 0.96578718 | 0.3 | 0.63894989 | 1.20447356 |
| 3 | 0.68331135 | 0.92571359 | 0.40356018 | 1.20420227 | 1.35024631 | 1.26211361 |
| 4 | 0.96146344 | 0.37275673 | 0.96610808 | 0.75038257 | 1.26650438 | 1.29541963 |
| 5 | 0.69223035 | 0.97243412 | 1.17496324 | 0.73516754 | 1.24553979 | 1.35627194 |
| 6 | 0.55066724 | 0.96347845 | 0.10820526 | 1.06598844 | 1.29776714 | 1.27910611 |
| 7 | 1.11262124 | 0.65023927 | 0.64227275 | 1.33353816 | 1.3162349 | 1.29989319 |
| 8 | 0.78234633 | 0.32142857 | 0.62534911 | 0.0830075 | 1.34890222 | 1.22927817 |
| 9 | 0.85056211 | 0.60949718 | 0.40645792 | 1.2428491 | 1.06745604 | 1.25850872 |
| 10 | 0.97209214 | 0.78990066 | 0.89551763 | 1 | 1.19063932 | 1.31476819 |
| 11 | 0.69328098 | 0.09441166 | 0.3050203 | 0.29657843 | 1.29025241 | 0.74968538 |
| 12 | 0.28188874 | 0.9669731 | 1.22090362 | 0.1169925 | 1.35733656 | 1.27015772 |
| 13 | 1.06924641 | 0.93169468 | 0.03571429 | 0.14150375 | 1.27256068 | 1.34576374 |
| 14 | 0.83532878 | 0.86647011 | 0.59787896 | 0.53219281 | 1.00358289 | 1.32127623 |
| 15 | 0.90082758 | 0.95006167 | 0.90651607 | 1.15065555 | 1.24181886 | 1.3181352 |
| 16 | 1.12741936 | 1.16930541 | 0.38410313 | 1.09248125 | 1.34462084 | 1.38411497 |
| 17 | 0.94026379 | 1.01658765 | 0.16374866 | 1.03257973 | 1.11478234 | 1.09955316 |
| 18 | 1.22249553 | 1.03685187 | 1.13752093 | 0.99068906 | 1.24553253 | 1.51453513 |
| 19 | 0.70769319 | 0.99006821 | 0.52363493 | 1.11065629 | 1.25292585 | 1.14394413 |
| 20 | 0.3090178 | 0.57221968 | 0.69534322 | 0.95216004 | 1.20452535 | 0.80808175 |
| 21 | 0.20838377 | 1.06415693 | 0.74422446 | 1.25041672 | 1.3004986 | 0.35686687 |
| 22 | 1.04414208 | 0.71627403 | 0.10714286 | 1 | 1.20976265 | 1.34431378 |
| 23 | 0.81863137 | 1.01558597 | 0.3187433 | 1.18371023 | 0.5992702 | 0.9137845 |
| 24 | 0.77428678 | 0.69809307 | 0.98152095 | 0.79366379 | 1.19494849 | 1.38846475 |
| 25 | 0.95431981 | 0.61747098 | 1.14700834 | 0.37598882 | 1.11092661 | 1.49837062 |
| 26 | 1.06200509 | 1.05243698 | 1.04377307 | 0.1169925 | 0.11321161 | 1.45951409 |
| 27 | 1.17748589 | 0.85243595 | 1.04125522 | 0.35108335 | 1.21091295 | 1.26717622 |
| 28 | 1.02449193 | 0.9846867 | 0.35633143 | 0.54431378 | 1.23501737 | 1.43353816 |
| 29 | 0.70446769 | 0.82035446 | 0.56899428 | 0.19967234 | 1.23319832 | 0.45395466 |
| 30 | 0.65930187 | 0.68832903 | 1.04866898 | 0.99772799 | 1.32590857 | 0.2 |
| 31 | 0.8158403 | 0.55060698 | 0.33996806 | 0.15849625 | 0.8263962 | 0.62407913 |
| 32 | 0.92821544 | 1.03767951 | 1.20698559 | 0.5 | 0.11595502 | 1.50891297 |
| 33 | 0.90263343 | 0.95049219 | 0.89434196 | 0.2299685 | 1.23633227 | 0.62419087 |
| 34 | 0.74990959 | 0.40670889 | 0.46312742 | 0.9194603 | 1.07652077 | 1.37139771 |
| 35 | 1.30573712 | 0.86764158 | 0.98416201 | 0.1169925 | 1.29667389 | 1.21301415 |
| 36 | 0.57407648 | 1.00402174 | 0.77471437 | 1.19858419 | 0.97046398 | 1.39163564 |
| 37 | 0.69373464 | 1.08973541 | 1.26907276 | 1.06438562 | 1.30892878 | 1.42041182 |
| 38 | 0.86274954 | 0.57086811 | 0.37654 | 1.23151496 | 1.30859543 | 1.28812369 |
| 39 | 1.11052423 | 0.19070893 | 0.80118862 | 1.1019417 | 1.11123277 | 1.30429845 |
| 40 | 1.16363931 | 0.68239273 | 0.73597283 | 0.59188632 | 1.34714326 | 1.24576374 |
| 41 | 0.31025775 | 0.70440812 | 1.17714937 | 1.09858419 | 1.30414223 | 1.06800298 |
| 42 | 1.09899887 | 1.03028933 | 0.93832419 | 0.9509775 | 0.83086675 | 1.30887882 |
| 43 | 1.02120064 | 0.77007585 | 0.9203289 | 0.40808175 | 1.33986971 | 1.53174597 |
| 44 | 1.06970496 | 0.75111087 | 0.87874437 | 0.2 | 0.31939515 | 1.49887069 |
| 45 | 0.41965783 | 0.75696406 | 0.7838964 | 1.12753486 | 1.20365572 | 0.38846475 |
| 46 | 0.68318868 | 1.00157957 | 0.94014636 | 0.5169925 | 1.30990997 | 1.42316337 |
| 47 | 1.11510448 | 0.97776 | 0.51172694 | 0.75038257 | 1.18687715 | 1.21357093 |
| 48 | 0.35658708 | 0.78473438 | 0.9950518 | 0.3 | 1.20095467 | 0.44576374 |
| 49 | 0.5841186 | 0.96386567 | 0.32020658 | 0.26767629 | 1.34738565 | 1.2258566 |
| 50 | 0.6865186 | 0.26844754 | 0.21428571 | 0.933985 | 0.94737447 | 0.67234 |
| 51 | 0.99740186 | 0.23728058 | 0.02696027 | 0.8 | 1.14735149 | 0.9950518 |
| 52 | 0.2343992 | 1.26351184 | 1.04299544 | 0.8 | 0.78302047 | 1.15481502 |
| 53 | 0.16042332 | 0.78684039 | 1.11587849 | 0.99083926 | 1.12030475 | 0.34330054 |
| 54 | 0.81002746 | 0.60953941 | 0.21428571 | 0.81032878 | 0.2947848 | ---- |
| 55 | 0.28893487 | 1.07336619 | 0.14285714 | 0.90947375 | ---- | ---- |
| 56 | ---- | 0.99543854 | 0.95528537 | 0.96232719 | ---- | ---- |
| 57 | ---- | 0.81517588 | 0.10714286 | 0.3 | ---- | ---- |
| 58 | ---- | 1.26762588 | 0.92306393 | 1.06219946 | ---- | ---- |
| 59 | ---- | 0.60449721 | 0.94750032 | 0.88137812 | ---- | ---- |
| 60 | ---- | 0.9382225 | 0.19946295 | 1.11357093 | ---- | ---- |
| 61 | ---- | 0.68801814 | 0.19946295 | 0.96211361 | ---- | ---- |
| 62 | ---- | 1.00723175 | 0.21428571 | 0.04150375 | ---- | ---- |
| 63 | ---- | 0.73484628 | 0.33963482 | 1.26800298 | ---- | ---- |
| 64 | ---- | 0.62672214 | 0.67506152 | 0.51083347 | ---- | ---- |
| 65 | ---- | 0.82513271 | 0.36921685 | 0.05849625 | ---- | ---- |
| 66 | ---- | 0.85640135 | ---- | 0.0509775 | ---- | ---- |
| 67 | ---- | 0.94903135 | ---- | 0.2 | ---- | ---- |
| 68 | ---- | 0.78134801 | ---- | 0.12288187 | ---- | ---- |
| 69 | ---- | 0.09838884 | ---- | 0.82288187 | ---- | ---- |
| 70 | ---- | 0.80426275 | ---- | 0.76617781 | ---- | ---- |
| 71 | ---- | 0.26280965 | ---- | 0.3 | ---- | ---- |
| 72 | ---- | ---- | ---- | 0.69858419 | ---- | ---- |
| 73 | ---- | ---- | ---- | 0.56438562 | ---- | ---- |
| 74 | ---- | ---- | ---- | 0.51083347 | ---- | ---- |
| 75 | ---- | ---- | ---- | 0.69645165 | ---- | ---- |
| 76 | ---- | ---- | ---- | 0.39138389 | ---- | ---- |

